# Methyl-TROSY NMR at 1.2 GHz: exploiting resolution and magnetic field alignment for structure and dynamics of large biomolecular assemblies

**DOI:** 10.64898/2026.10.21.760675

**Authors:** Vincenzo R. Lobbia, Andrea Orsetti, Hugo van Ingen

## Abstract

The range of protein assemblies that can be studied by NMR has greatly expanded both by increases in magnetic field strength and by development of new NMR approaches, such as methyl-group specific transverse relaxation optimized spectroscopy (methyl-TROSY). Recently, a new generation of NMR magnets operating at 1.2 GHz have become available, offering the highest resolution for biomolecular NMR. Here, we investigated the performance of methyl-TROSY NMR at 1.2 GHz in the characterization of the structure and dynamics of large biomolecular assemblies, using the nucleosome, a 200 kDa protein-DNA complex, as a test sample.

We find that the increased resolution of the 1.2 GHz system allowed to resolve small asymmetries in sidechain conformations that remained hidden at lower magnetic field strengths. Increased CSA relaxation effectively increased ^13^C transverse relaxation rates by 20% at 1200 compared to 900 MHz, negatively impacting sensitivity. We further observed significant magnetic field alignment of the nucleosome at 1.2 GHz, giving rise to methyl ^1^H-^13^C residual dipolar couplings (RDCs). We show that these histone methyl group RDCs can be used to aid stereo-specific assignment and to determine the overall conformation of the nucleosomal DNA, revealing a significant unwrapping of the DNA from the histone core in solution. We expect that 1.2 GHz NMR will be very advantageous for studies of large biomolecular assemblies, and in particular for protein-DNA complexes.

## Introduction

With availability of higher magnetic fields, the increased resolution and sensitivity have enabled NMR spectroscopists to study more complex systems. Next to these instrumental factors, the development of new labeling approaches and transverse-relaxation optimized spectroscopy (TROSY), and in particular methyl-TROSY, have greatly expanded the size limit of the system that can be studied using solution NMR^1–10^. Thanks to development of high-temperature superconducting materials, the leading commercial manufacturer in NMR spectrometers has been able to build a high-resolution NMR machine operating at 28.2 T or 1H resonance frequency of 1.2 GHz. The performance of solution NMR at this ultra-high field has been discussed in a few recent papers^11,12^.

As anticipated early on^65^, the increased magnetic field will also increase the effects of magnetic field alignment. When molecules with an anisotropic magnetic susceptibility are placed in a magnetic field, the induced magnetic moment will depend on their orientation with respect to the field, causing them to align in such a way that the induced magnetic moments are minimal^13,14^. Nucleic acids are well known to have a large magnetic susceptibility anisotropy (χ_a_) due to the nearly colinear arrangement of the aromatic bases, and thus show pronounced magnetic field alignment^13,15,16^. The net alignment of the molecule causes incomplete averaging of the dipolar interactions and hence result in residual dipolar couplings (RDCs) which can be used to determine structure^17–20^. Proteins show less magnetic field alignment compared to nucleic acids as they in general have few aromatic residues which will be not colinear. Yet, even for ubiquitin at moderate magnetic field strength of 750 MHz small magnetic field induced RDCs (fiRDCs) have been measured.^21^ As the alignment depends on the square of the magnetic field, significant fiRDCs for both proteins and nucleic acids can be expected at 1.2 GHz. These fiRDCs will add to the regular J-coupling splitting and “traditional” RDC splittings when using anisotropic solutions, as has been evaluated in detail for nucleic acids^22^. Importantly, as magnetic susceptibility anisotropy can be predicted quite accurately from the structure of a molecule^23^, fiRDCs can be used for structure determination^17–20,24^.

Fo large biomolecular assemblies, the symmetric stacking of subunits can increase the magnetic susceptibility anisotropy of the assembly, increasing the field-induced alignment. Thus, methyl-TROSY NMR at 1.2 GHz also offers exciting new potential for structure determination of large biomolecular assemblies, in particular for large complexes involving nucleic acids, such as the nucleosome.

Here, we compared methyl-TROSY spectra recorded at 600, 900 and 1200 MHz in terms of sensitivity, resolution and extracted fiRDCs. We found that methyl-TROSY spectra at the 1200 MHz system were ∼20% less sensitive than at 900 MHz system, likely due to a combination of increased transverse relaxation and hardware and sample factors. At 1200 MHz, resonances for several isoleucine, leucine and valine (ILV) methyl groups show clearly resolved splittings that correlate with asymmetric rotamer states in many nucleosome crystal structures. Further, we observe large fiRDCs of up to 8 Hz at 1200 MHz. We show that these can be used to obtain stereospecific assignments of nearly all methyl groups. Importantly, the experimental fiRDCs can be used to determine the conformation of the linker DNA in the nucleosome, revealing significant unwrapping of the nucleosomal DNA from the histone core.

## Materials and Methods

### Expression and purification of isotope labelled histones

The histones, H2A, H2B, H3 and H4, from *Xenopus laevis* (*Xl.)* were transformed into BL21(DE3) pLysS cells using pET3a derived plasmids encoding histones (Uniprot P06897, P02281, P84233, P62799). Histones were perdeuterated using an adapted protocol of Marley et al^30^. In short, cells were grown at 37 °C in Luria-Bertani (LB) medium to optical density (OD) 1.5, then spun down and washed twice in D_2_O-based M9 medium without glucose and NH_4_Cl. Cell pellets were resuspended in D_2_O M9 with deuterated glucose, and ^15^NH_4_Cl for labeled histones or ^14^NH_4_Cl for fully deuterated histones to an OD of ∼0.6 and incubated for 1–2 hours at 37 °C. To obtain perdeuterated ^15^N,{^1^H-^13^C}-ILV labeled H2B, 60 mL/L α-ketobutyric acid (4-^13^C,3,3-2d) and 80 mg/L α-ketoisovalerate (3-(methyl-d3),4-^13^C,3-d) sodium salt (Sigma-Aldrich) were added to the culture and incubated for another hour, while for H2B with {^1^H-^13^C}-labeling of both LV methyl groups (2MeLV), 80 mg/L 2-Keto-(3-methyl-^13^C)-butyric-4-^13^C,3d acid (Sigma-Aldrich) was added. All histones were expressed overnight with 1 mM IPTG at 37 °C.

Histones were purified as described previously in Klinker et al.^31^. Briefly, cell pellets were resuspended in lysis buffer (50 mM Tris, 100 mM NaCl, 1% triton, pH 7.5) with lysozyme, then lysed using one freeze-thaw cycle and sonication. The inclusion bodies were collected and unfolded using unfolding buffer (6 M Guanidinium-HCl, 50 mM Tris, 100 mM NaCl, pH 7.5). Unfolded histones were purified using first size-exclusion chromatography (SEC) over a superdex 75 column (GE Healthcare) in denaturing buffer (7 M urea, 50 mM KPi, 1 mM EDTA, 150 mM NaCl, pH 7.5) and then using ion-exchange chromatography (IEX) over a combined HiTrap Q and HiTrap SP column (Cytiva) in denaturing buffer, where the HiTrap Q column was removed before eluting the histones from the SP column using a salt gradient (0.15 – 1 M NaCl in denaturing buffer). Fractions were analyzed using SDS-PAGE, histone containing fractions were pooled and concentrated to 3 mg/mL, supplemented with 1 mM lysine, and stored at -20°C for further use.

### DNA production

A pUC19 plasmid containing 12 x 167 base pair repeats was used to produce 167 base pair 601 Widom DNA^32^. Plasmids were produced using DH5alpha in 2xYT medium with an overnight culture at 37 °C. Plasmids were isolated using alkaline lysis^33^ followed by isopropanol and ethanol precipitation and redissolved in TE buffer (10 mM Tris, 1 mM EDTA, pH 8.0). Plasmids were purified using IEX over HiTrap Q column (Cytiva) with a 0-1 M NaCl gradient in TE buffer. Purified plasmids were digested in TE buffer supplemented with 10x Cutsmart buffer (New England Biolabs) to 1x final concentration using 700U ScaI-HF (New England Biolabs) per 10 mg plasmid DNA for 72 hours at 1mg/mL plasmid concentration. Completion of the restriction was monitored using PAGE. Then 167 bp DNA fragments were purified using anion exchange (HiTrap Q column 5 mL), ethanol precipitated and resuspended in TE buffer to a concentration of ∼2.4 mg/mL before storage at -20 °C.

### Histone octamer refolding and reconstitution of nucleosomes

The histone octamer was refolded as described in Dyer et al^33^. In brief, histones were mixed in equimolar ratio in denaturing buffer (7 M urea, 50 mM Tris, 100 mM NaCl, pH 7.5) at 1 mg/mL protein concentration, using absorbance at 280 nm to determine their concentrations. The histone octamer mixture was dialyzed in three steps for 16, 8, and 16 hours to high salt buffer (2 M NaCl, 10 mM Tris, 1 mM EDTA) at 4 °C using 3 kDa MWCO membrane and then purified using SEC over a superdex 200 column (120 mL, GE Healthcare) in high salt buffer. Peak fractions were analyzed using SDS PAGE. The histone octamer eluted at 64 mL, with a yield of ∼70%. Fractions containing pure octamers were pooled and concentrated to ∼4 mg/mL and stored at 4 °C before immediate use.

Nucleosomes were reconstituted from 601 DNA and perdeuterated histones and ILV-labeled H2B according to the protocol described by Dyer et al.^33^. Briefly, stock-solution of 601 DNA was adjusted with 4 M NaCl to a final salt concentration 2 M and ∼3 mg/mL DNA. Equimolar amounts of histone octamer (in high salt buffer) and 601 DNA were mixed at 4 °C and diluted to 0.7 mg/mL protein concentration using high salt buffer, incubated for 2 hours, and then dialyzed using a 16-hour gradient to low salt buffer (0.25 M KCl, 10 mM Tris, pH 7.5, 1 mM EDTA). The soluble fractions containing nucleosomes (ca. 70% yield based on DNA absorbance) was then buffer exchanged to NMR buffer (20 mM KPi, pH 6.2, 0.01% NaN_3_, 100% D_2_O) and concentrated to ∼13.6 mg/mL over a 30kDa MWCO centrifugal concentration device. Analysis by SDS-PAGE and native PAGE confirmed that the obtained nucleosomes are free of significant amounts of free DNA (Supplementary Figure S1). Nucleosomes were stored at 4 °C before use.

### NMR spectroscopy

All NMR experiments were recorded on Bruker Avance spectrometers, using either an Avance III 600 MHz spectrometer with 5 mm room-temperature probe, an Avance III 900 MHz spectrometer with a 5 mm cryo-probe, or an Avance NEO 1200 MHz spectrometer with a 3 mm cryo-probe. Samples were measured in a 5 mm Shigemi tube (600, 900 MHz) or a 3 mm tube (1200 MHz), in all cases containing 64.3 μM nucleosomes in 300 μL NMR buffer (20 mM KPi, pH 6.2, 0.01% NaN_3_, 100% D_2_O). All experiments were recorded at 45 °C. NMR data were processed using Bruker Topspin using forward linear prediction in the indirect dimension(s) and cosine-squared window functions, and subsequently analyzed using NMRFAM-SPARKY^34^, except where noted otherwise.

To compare resolution and sensitivity, methyl-TROSY HMQC experiments were recorded at 600, 900 and 1200 MHz. Experiments used the SOFAST fast-pulsing method^29^ with a PC9-shaped methyl selective pulse with 120° flip angle and Reburp 180° pulse (both with 6 ppm band width), a recycle delay of 0.5 s, and acquisition times of 66.5 ms (^1^H) and 42.4 ms (^13^C), and 3.3 kHz WALTZ64 decoupling during acquisition. Total measurement time was 2 hours for all three spectra. Peak intensities and noise levels were determined using the built-in functions of NMRFAM-SPARKY using data processed without linear prediction.

Sucrose measurements showed that the 1.2 GHz offers same or up to 30% more sensitivity compared to the 900 MHz system, depending on the ionic strength of the sample. Extrapolating from these measurements, one would expect ca. 10% sensitivity increase at 1.2 GHz at the low ionic strength (*I* = 25 mM) of the nucleosome sample. Accounting for differences in the inner diameter of the tubes used for the sucrose measurements and our nucleosome sample, which result in relative number of active spins of 33% for sucrose and 28% for the nucleosome at 1.2 GHz, we expected a relative sensitivity at 1.2 GHz of ∼93% (0.28/0.33*1.1).

### Resonance assignment Xl. H2B ILV methyl groups

Initial assignments of the ILV methyl group resonances in *Xenopus leaevis (Xl.)* H2B were based on assignment transfer from the available *Drosophila melanogaster* (*Dm.*) chemical shifts^35^. To confirm and complete the assignments, 3D methyl-TROSY (h)CCH NOESY were recorded with at 900 (600) MHz and acquisition times *t*_1,max_ 8 ms, *t*_2,max_ 8 ms and *t*_3,max_ 67 ms, total measurement time 3 days, and with 200 (50) ms NOESY mixing time on nucleosome sample with ILV (2MeLV)-labeled H2B. Assignments were made using MAGMA 1.2.3^36^ and verified manually by comparing the experimental NOE network and to the methyl-methyl distances in the 1KX5 nucleosome crystal structure^37^. Stereospecific assignments could be transferred from the *Dm*. spectra based on conservation of NOE patterns for 16 of the 30 LV resonances (5 out of 6 Leu and 3 out of 9 Val residues). Comparison of the Leu ^13^C δ1 and δ2 chemical shifts and the crystal structure rotamer conformation using the Sider program^38^ confirmed the stereospecific assignments for all Leu residues, except L98 and L99, which were too dynamic.

### ILV rotamer analysis

Nucleosome crystal and cryo-EM structures in the PDB were selected if they contained *Xl.* canonical histones and had a resolution below or equal to 3.5 Å. The side chain conformation of ILV residues was analyzed to determine their rotamer distributions. Twenty nucleosome structures that contained all eight *Xl.* histones, two DNA strands and no other binding partners (excluding ions) were selected (see Table S1). After adapting the H2B residue numbers for consistency, the H2B histone chains were extracted using PyMOL and subsequently all H2B structures were aligned on the backbone N, CA, CO atoms of each ILV residue, one at a time.

### Sensitivity measurements

Sensitivity curves for SOFAST methyl-TROSY experiments^29^ were measured from 2D spectra recorded with *t*_1,max_ of 15 and *t*_2,max_ of 62 ms and recycle delay values ranging from 0.2 to 4 sec using ^1^H shaped pulses with either 2 ppm or 6 ppm bandwidth at 600, 900 and 1200 MHz. Peak intensities (*I*) were measured in NMRFAM-Sparky (ref) and converted to sensitivity-per-unit-time (S/N)*_t_* by normalizing the peak intensities to the measurement time and using the dependence of the noise level per unit of time on the total scan time Tscan (refs):

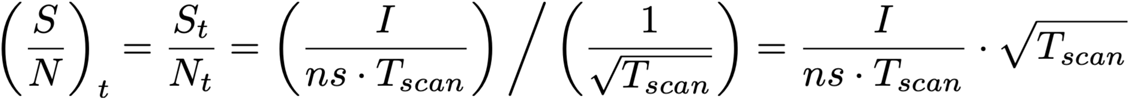

where *ns* is the number of scans, *T*_scan_ equals *d1* + *aq* + *t*_seq_, where *d1* is the recycle delay, *aq* the ^1^H direct acquisition time and *t*_seq_ is the total average time of the sequence accounting for all delays and average *t*_1_ acquisition time of the pulse sequence (16 ms). These (relative) sensitivity curves were fit to:

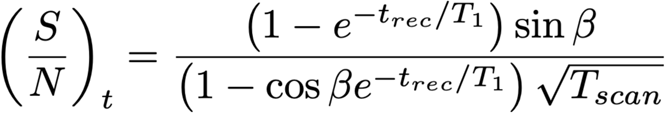

to extract the optimal *T*_scan_ per residue and per flip-angle β.

### Relaxation measurements

The longitudinal relaxation rates (*R*_1_) of methyl group ^1^H spins were measured by implementing a saturation recovery element using either a 90° hard uniform or a 90° PC9-shaped selective excitation pulse (band width 6 ppm) before the methyl-TROSY experiment^39^. Spectra were recorded with an inter-scan delay of 6 sec and relaxation delay values of 50, 100, 150, 200, 800, 1200 and 3000 ms.

The transverse relaxation rates (*R*_2_) of methyl group ^13^C spins were measured using a ^13^C-^1^H MQ Carr–Purcell–Meiboom–Gill (CPMG) relaxation dispersion experiment^40^ with or without a constant CPMG relaxation time T_relax_ of 10 ms and CPMG pulsing rates ν_CPMG_ of 100 and 1000 Hz. Values at highest pulsing rates were used for analysis (1000 Hz). MQ data were recorded with offset set to center of Ile and Leu/Val regions to minimize off-resonance effects.

Both datasets were processed using NMRPipe^41^ with exponential line broadening and peak volumes were obtained by fitting the 2D peaks to a gaussian line shape in the program PINT^42^. To obtain the methyl ^1^H *R*_1_ rates, peak volumes were fit to Eq. [1]:

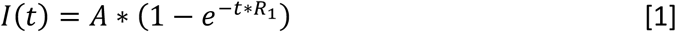

where *I*(*t*) is the peak volume at relaxation delay *t*, *A* is a scaling factor. The methyl ^13^C effective *R*_2_ rates, *R*_2,*eff*_, were calculated using Eq. 2:

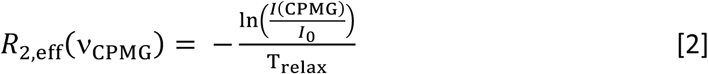

where *I*_0_ is the peak volume in the reference spectrum and *I*(ν_CPMG_) the peak volume obtained in the CPMG experiment with pulsing rate ν_CPMG_. Error bars in the fitted relaxation rates were calculated using PINT, which were estimated using the jackknife method.

### RDC measurement

The ^1^H-^13^C one-bond methyl fiRDCs (*D*) were derived from methyl-TROSY IPAP spectra that encode the *J* + *D* peak splittings in the ^1^H dimension^43^. Data were recorded at 600, 900 and 1200 MHz with FID resolution of 7 Hz and processed to spectra with digital resolution of 2.35 Hz (typical line width in the ^1^H dimension was ∼30 Hz). Spectra with either the inphase (IP) and antiphase (AP) doublets were added/subtracted to result in spectra with either the upfield or downfield doublet component. Peak positions of the upfield and downfield component were determined using the ‘peak center’ command in NMRFAM-SPARKY and then subtracted as values in Hz to yield the peak splitting (Δ). The error in Δ (σ_Δ_) was derived from the error in peak position using the error propagation (Table S2). The error in peak position was estimated to be LWHH/(S/N), where LWHH is the linewidth at half height and S/N the signal-to-noise ratio. Peaks with severe overlap (V38a, V108γ1, L42δ2a and L42δ2b) were excluded from the analysis. The fiRDCs were obtained by fitting peak splittings Δ using gnuplot 5.4^44^ to Eq. [3]:

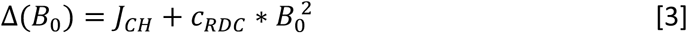

where *J*_CH_ is the one-bond 1H-13C scalar coupling constant, *c*_RDC_\**B*_0_^2^ is the fiRDC at magnetic field strength *B*_0_, and *c*_RDC_ is a scaling coefficient that depends, amongst others, on the magnitude of the magnetic susceptibility anisotropy and the bond-vector orientation. The fiRDC at 1200 MHz was calculated as *c*_RDC_\**B*_0_^2^, where *B*_0_ is 28.2 T, and the error was set to at least 0.5 Hz or σ_cRDC_\**B*_0_^2^, where σ_cRDC_ is the fit error on *c*_RDC_ (Table S3).

### Determination of magnetic susceptibility tensor from experimental fiRDCs

The experimental fiRDCs at 1200 MHz were used to determine the magnitude and orientation of the alignment tensor using the DC program of NMRPipe as implemented on the webserver (https://spin.niddk.nih.gov/bax/nmrserver/dc/svd.html) in the following way. First, only the fiRDCs from resonances with known stereo-specific assignment without overlap (20 in total) were used to fit alignment tensor, using the H2B structure (single chain) from 1KX5^37^, 3LZ0^45^ or 7OHC^46^. Error on the fiRDCS was set to 1 Hz for all residues. Comparison of observed vs. calculated fiRDCs identified three outliers (from I58 and L98 methyl groups) in the fits using the 1KX5 and 7OHC structures. After removing these outliers, the best fit according to Q-factor was obtained for the 1KX5 structure. The calculated RDCs from this fit were used to assign the stereochemistry of the 6 LV residues (excluding V15 in the histone tail) without stereospecific assignment. For each of these residues, the measured RDCs were compared with the predicted RDCs for the two possible assignments, evaluating for each assignment possibility Eq. 4:

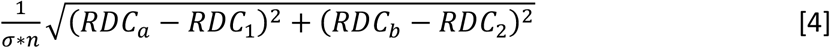

where RDC_a_ and RDC_b_ are the observed fiRDC for the two methyl groups of the same residue, RDC_1_ and RDC_2_ the predicted RDCs, *n* the amount of measured RDCs per residue (n = 1 if RDCs are available for only one methyl group due to overlap or low S/N), and σ is the standard deviation of the measured set of RDCs. Confident assignments were made when the score of equation 4 was less than 4 and the difference between the score of the two possible assignments was more than one σ, resulting in stereospecific assignments for all 6 remaining residues. Eq. 4 was similarly used to determine stereospecific assignments when including only fiRDCs from Ile or Ile and Sider-predicted Leu in the fit.

The final list of 29 fiRDCs was used to calculate the alignment tensor using 1KX5. The final magnetic susceptibility tensor was calculated from the fitted alignment tensor by scaling the eigenvalues with *B*_0_^2^/15*kT*, corresponding to 82814 at 28.2 T and 45 °C. The error in the magnetic susceptibility tensor was estimated by adding noise to the residual dipolar couplings, or noise to the reference structure, and by random removal of residual dipolar couplings from the dataset using the default “advanced error analysis options” on the webserver^47^. The final reported error is calculated as the average error in the different methods.

### Prediction of magnetic susceptibility tensor from structure

To calculate the magnetic susceptibility tensor from known 3D structures, we followed the approach of Bryce et al.^23^, taking into account the contributions from both DNA and protein. The residue-specific magnetic susceptibility anisotropies for DNA bases and protein carbonyl groups and aromatic residues were set to their axial symmetric values and orientations as described in Bryce et al. The residue-specific contributions were added via tensor-summation to the overall molecular magnetic susceptibility anisotropy.

To determine the linker DNA conformation in the 167 bp 601-nucleosome, structural models were constructed by extending the 147 bp 601-nucleosome crystal structure (3LZ0) with linear B-DNA that was generated using the 3DNA webserver^48^. To generate models of 167 nucleosome with different conformations of the linker DNA, the linear DNA fragment was added to the 3LZ0 DNA at bp ±70 (closed linkers), ±60 (straight linkers), ±55 (open), and at bp ±70 for one arm and bp ±55 for the other arm (asymmetric linkers).

## Results

For our study we used nucleosomes reconstituted in vitro from perdeuterated *Xenopus laevis (Xl.)* histones with ILV-labeled histone H2B and a 167 bp DNA fragment corresponding to the Widom 601 strong positioning sequence^32^. Reconstitution of the *Xl.* 601-nucleosome was efficient and yielded pure samples (Fig. S1) that gave highly resolved 2D ^1^H^13^C methyl-TROSY NMR spectra (Fig. S2) with the expected 36 cross peaks for each of the H2B Ile δ1, Leu δ1/δ2, and Val γ1/γ2 methyl groups. Peak assignments were transferred from the previous *Drosophila* chemical shifts assignments^35^ where possible, resulting in the confident assignment of 27 methyl group signals. The assignments were completed through analysis of 3D NOESY experiments using the known nucleosome structure^37^. The transferred assignments and NOESY data also allowed to determine the stereospecific assignment of methyl groups in 5 out of 6 Leu and 3 out of 9 Val residues.

### Resolution and sensitivity of methyl-TROSY NMR at 1.2 GHz

We recorded methyl-TROSY spectra at 600, 900 and 1200 MHz and evaluated the obtained resolution and sensitivity. As the methyl-TROSY effect relies on relaxation interference between intra-methyl ^1^H-^13^C dipolar interactions it is purely dipolar in nature and field independent^6,25^. However, the ^13^C spin in a methyl group has a small, but non negligible chemical shift anisotropy (CSA)^26^, causing a field-dependent increase to the transverse relaxation rate^27^.

Spectral comparison shows clearly that peak overlap that is present at 600 or 900 MHz is resolved at 1200 MHz (Fig. 1a and S2). More detailed analysis shows that when expressed in ppm, the line widths at 1200 MHz are decreased by 23% in the ^1^H and 7% in the ^13^C dimension compared to the line widths at 900 MHz. While the resolution increase in the ^1^H dimension is close to that expected from the increase in magnetic field, the resolution increase in the ^13^C dimension is significantly less, due to an increase in ^13^C line widths at 1200 MHz (Fig. S3).

**Figure 1:**
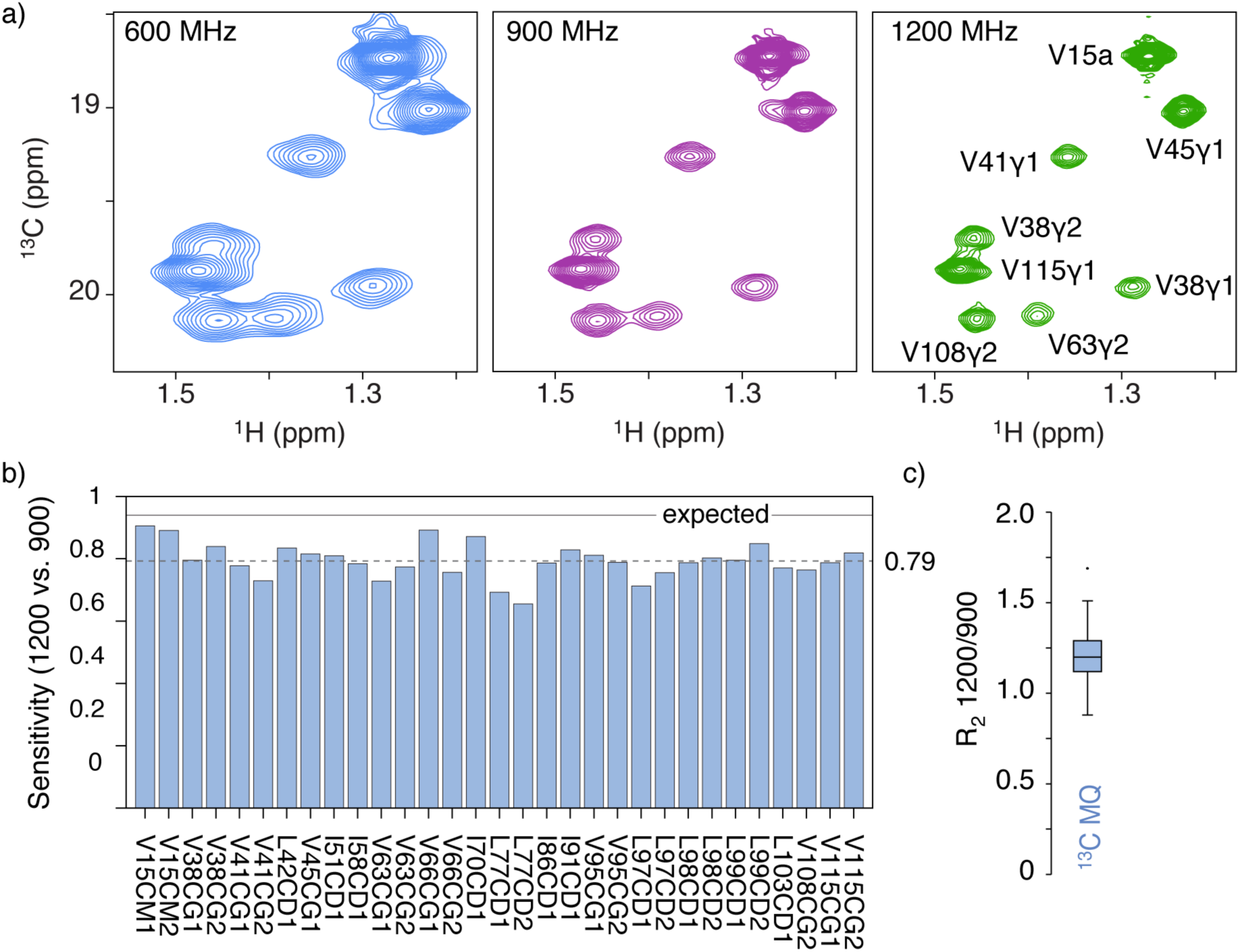
Resolution and sensitivity of methyl-TROSY NMR at 1.2 GHz. a) Zoom of methyl-TROSY spectra of nucleosomes with ILV-labeled H2B at indicated magnetic field strengths highlighting the increased resolution at 1.2 GHz. Assignments are shown on the 1.2 GHz spectrum only. Assignment V15a is not stereospecific. b) Relative sensitivity of methyl-TROSY signals of H2B ILV residues in the nucleosome at 1200 MHz in a 3 mm tube compared to 900 MHz in a 5 mm tube at constant concentration; solid line: expected sensitivity as extrapolated from the sucrose sensitivity standards and considering ionic strength and sample volumes; dashed line: average sensitivity for all methyl signals. c) Boxplot of the ^13^C-^1^H MQ R_2_ rates at 1200 MHz compared to 900 MHz, showing on average 1.2x increase.

To compare the sensitivity of the methyl-TROSY spectra, we focused on the 900 and 1200 MHz spectra as these were both recorded using cryo-probes. Notably, the 1.2 GHz system is equipped with a 3 mm probe, whereas the 900 MHz system has a 5 mm probe. As we used the nucleosome sample at the same concentration, the number of spins in the active volume of the 1.2 GHz is ca. 28% of that for the 900 MHz. Despite this, the sensitivity at 1200 MHz is close to that of the 900 MHz system, albeit on average 20% lower (Fig. 1b). To explain this reduction in sensitivity, we measured ^1^H and ^1^H-^13^C multiple-quantum (MQ) *R*_2_ relaxation rates. On average, the ^1^H-^13^C MQ *R*_2_ rates are 1.2 times higher at 1200 vs. 900 MHz (Fig. 1c), while there was no notable change in ^1^H R_2_ (Fig. S3). This is in-line with the expected impact of the CSA contribution to the transverse relaxation of methyl groups^27^. Studies on ubiquitin have shown that the ^1^H CSA is very small, on the order of 1 ppm, while the ^13^C CSA is sizable, with average value of 18 ppm for Leu/Val and 25 ppm for Ile methyl groups^26^. The increased ^13^C CSA relaxation at 1.2 GHz thus explains most of the observed sensitivity loss.

The reduction in sensitivity could additionally be caused by differences in steady-state ^1^H magnetization as we recorded these experiments using the SOFAST fast-pulsing approach^28,29^. In this approach the sensitivity gain depends on the longitudinal relaxation rate of the ^1^H spins, which will decrease slightly with increased magnetic field. To investigate this we measured methyl group ^1^H longitudinal relaxation times (*R*_1_) rates of the H2B ILV methyl groups in the nucleosome. We used both a methyl-selective (R_1,sel_) and a non-selective ^1^H excitation pulse (R_1,uni_) at 900 MHz and then compared the R_1,sel_ different magnetic field strengths (Fig. 2a-c). Overall, the ^1^H R_1_ relaxation is more efficient when using the selective pulse, indicating a significant ‘spin cooling’ effect by keeping non-methyl protons in equilibrium (Fig. 2a, b). The I36 methyl group experiences the largest increase in relaxation rate which can be explained by its proximity to the protonated DNA. The other H2B methyl groups are however far from the DNA. For instance, V63 and V66 experience a large increase in relaxation rate but are in the core of the nucleosome at considerable distance from the DNA (12.5 and 19 Å, respectively). This indicates that enhancement of R_1_ relaxation for the nucleosome core methyl groups is likely due to the presence of non-methyl ^1^H spins within the histone and/or very efficient spin-diffusion throughout the histone ^1^H-^1^H network to the protonated DNA. Comparison of R_1,sel_ rates at 600, 900 and 1200 MHz show a negligible increase with magnetic field (Fig. 2c), indicating that the degree of methyl-group polarization recovery during fast-pulsing experiments should be maintained at 1.2 GHz.

**Figure 2:**
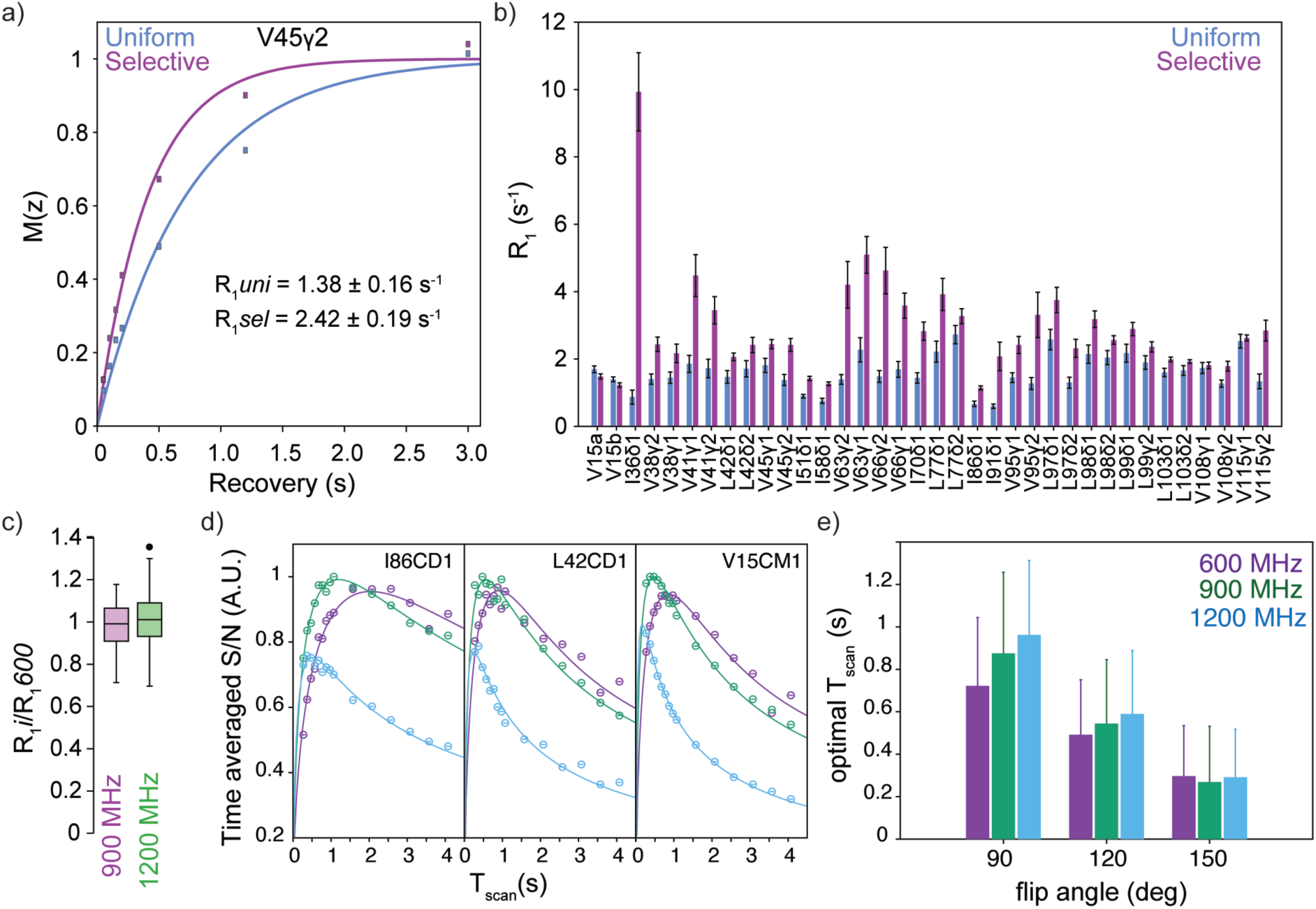
Methyl group 1H longitudinal relaxation rates and optimal flip-angle and scan time for SOFAST methyl-TROSY. a) Saturation recovery curve of V45γ2 methyl together with best fit and fitted 1H *R*_1_ values upon either methyl-group selective or uniform excitation. b) Comparison of ^1^H *R*_1_ values obtained after methyl-selective (*R*_1,sel_) or uniform excitation (*R*_1,uni_) for all H2B ILV residues at 900 MHz. Most core residues have significantly larger R_1,sel_. c) Boxplot of R_1,sel_ values at 900 or 1200 MHz compared to 600 MHz. The outlier is V38γ2. d) Sensitivity of the SOFAST methyl-TROSY experiment as function of total scan time T_scan_ for methyl-selective excitation with 90 (purple), 120 (green) or 150° (blue) pulse, all at 1200 MHz. Solid lines show the best-fit of the data to expected sensitivity curves for SOFAST experiments^49^ and was used to extract the optimum T_scan_. e) Comparison of the optimal total scan time T_scan_ at 600, 900 and 1200 MHz when using a 90, 120 or 150 excitation pulse.

We also compared the sensitivity and optimal total scan time (T_scan_) of the SOFAST methyl-TROSY experiment using three different pulse angles at 600, 900 and 1200 MHz (Fig. 2d, e). Optimal signal-to-noise was obtained using a 120° selective excitation pulse at all fields. The optimal T_scan_ time increases slightly with increasing field, in particular when using a 90° selective excitation pulse. Such field-dependent optimal inter-scan delay has also been observed for SOFAST ^1^H,^15^N-HSQC experiments^49^. We thus find that the optimal T_scan_ increases from 0.45 s at 600 MHz to 0.55 s at 1200 MHz. At the constant T_scan_ (0.5 s) used in the sensitivity comparison of Figure 1b the slight mis-setting of optimal T_scan_ contributes less than 1% to the decrease in sensitivity at 1.2 GHz.

Finally, magnetic field induced alignment (see below) will contribute only modestly to reduced sensitivity. As the histones are perdeuterated no ^1^H-^1^H RDCs can form and sensitivity loss is limited to reduced transfer efficiency (∼1%) due to dispersion of the effective ^1^H-^13^C coupling values (J + *D*). We conclude that at 1.2 GHz the sensitivity of the methyl-TROSY experiment is reduced by the increased ^13^C CSA relaxation to the transverse relaxation and not significantly affected by changes in longitudinal relaxation when using the fast-pulsing approach. In addition, it is likely that hardware-related differences between the two systems also contribute to the sensitivity difference.

### Ultra-high field NMR visualizes heterogeneity in histone side chain conformations

Careful comparison of the recorded methyl-TROSY spectra showed to our surprise that three cross peaks that are very broad at 600 MHz are resolved into two sharper peaks at 1200 MHz with close to 1:1 intensity ratio (Fig. 3a). For residues I36 and V45, the two peaks have slightly different ^13^C chemical shifts, while for L103 the difference is in both ^1^H and ^13^C dimension. Previously, we observed peak splittings for residues in histones H3 and H2A that are in close contact with the nucleosomal DNA^35,50^. As the 601 DNA sequence we use is non-palindromic, residues in the DNA binding interface will experience two different chemical environments. This most likely also explains the double peak appearance for the H2B I36δ1 methyl group, which is close to the nucleosomal DNA.

**Figure 3:**
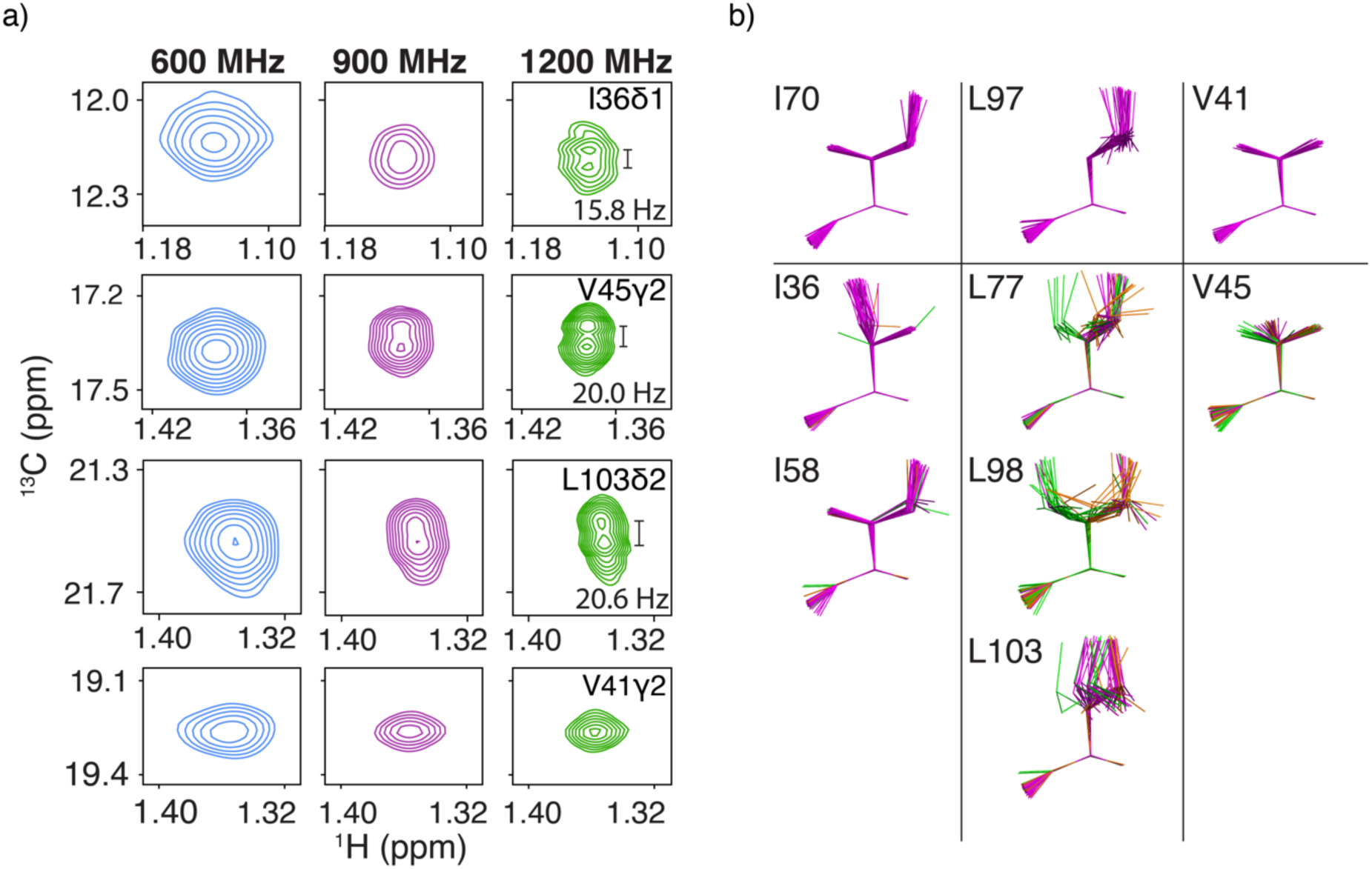
Asymmetry in the symmetric H2B histone copies of the nucleosome. a) Zoom on the peak shape for selected residues. At 1.2 GHz several residues I36δ1, V45γ2, L103δ2 show a split peak appearance, while most resonance such as V41γ2 have a normal peak shape. b) Heavy backbone atom superposition of selected Ile, Leu and Val side chains in 20 nucleosome crystal structures with Xl. histones (see Table S2). Side chains with difference in χ1/χ2 <= 30° between each H2B copy in a structure are color coded in magenta. Side chains for which the difference in χ1/χ2 > 30° are colored in green for one copy, and orange for the other copy.

For the other residues (V45 and L103), the peak splitting cannot be directly related to the DNA sequence as they are much further away. We can exclude that the splitting is caused by a *J*-coupling that is only resolved at 1200 MHz because 1) no *J*_CC_ couplings can be present due to the ILV labeling scheme; 2) *J*_CN_ couplings are only relevant for Val and are much smaller than the observed splittings (ca. 2 Hz^51^); 3) the splitting differs for the two residues; and 4) for L103 the split peaks are offset in both ^13^C and ^1^H dimension. Therefore, the split peak appearance indicates that the affected methyl groups experience subtly different chemical environments, unrelated to the DNA sequence. As peak splitting is on the order of 20 Hz (125 rad s^-1^), any interconversion between the two states occurs slowly, if at all, with approximate maximum exchange rate *k*_ex_ of 12 s^-1^.

Interestingly, comparison of the H2B ILV rotamers in twenty nucleosome structures containing *Xl.* histones show that, while most residues have only a single side chain conformation, a few have multiple conformations (Fig. 3b). The residues with multiple conformations in the crystal or cryo-EM structures match with the residues whose resonances either show peak splitting (V45, L103), have more peak broadening compared to other residues (I58, L77, L98), or show a skewed peak shape (L98, L103) at 1200 MHz (Fig. S4). Moreover, these residues have different conformations for each H2B copy in many of the structures analyzed. In 65% of the structures analyzed, residue V45 is in the gauche– conformation in one H2B copy but in the trans or an eclipsed conformation in the other copy. Also, for L103, L77 and L98, there are significant differences in the χ_1_ or χ_2_ dihedral angles between the two H2B copies in respectively, 40, 60 and 70% of the analyzed structures. For I58 only a single structure in our analysis shows a different side chain conformation between the two H2B copies, although its conformation is variable between different structures. Together, this suggests that the asymmetry seen in some crystal structures is also present in solution, resulting in different (distributions of) side chain conformation of these residues. Of note, the NMR data further indicate that the residues with asymmetric side chain conformations are also affected by a dynamic conformational exchange. This is evident from the slight shift in peak position between the different fields, which is most visible for I36δ1 (Fig. 3a). At this point we cannot tell the origin of this process and how it is related to asymmetry in sidechain conformations seen in the crystal and cyro-EM structures. A recent solid-state NMR study suggested that the histone octamer contains several dynamic hotspots in H3 and H4 that could be connected in a dynamic network that extends from the DNA binding surface into the octamer interior^52^. The data here suggest that such dynamic network may also be present within the H2A-H2B dimer, or at least within H2B, as the identified residues are similarly distributed from the DNA binding site to the interior.

### Substantial magnetic field induced alignment of nucleosomes at 28.2 T

We next examined the degree of magnetic field induced alignment of the nucleosomes. The expected molecular magnetic susceptibility anisotropy Δχ_mol_ was calculated for various model structures using residue-specific values of Δχ for the nucleobases, aromatic groups and the carbonyl backbone groups. To understand the alignment of the nucleosome, we first compared the Δχ_mol_ tensor for linear DNA and DNA arranged in a super-helix, as in the archaeal hyper-nucleosome^53,54^ (Fig. 4). For linear DNA where the bases are stacked colinearly, the individual Δχ tensors add up almost perfectly, resulting in Δχ_mol_ tensor with a main axis along the helical axis and thus a preferred molecular orientation orthogonal to the external magnetic field. When DNA is arranged in a super-helix, the main axis will be aligned with super helical axis (Fig. 4). As nucleosome core particle, which has 147 bp of DNA, has its DNA arranged in superhelix, we thus expect similar alignment with preferred orientation such that the superhelical axis is orthogonal to external magnetic field. Using the crystal structure of the nucleosome core particle (1KX5) and considering both DNA and protein contributions, the predicted Δχ_mol_ tensor has a main axial component of 105 10^−27^ J T^−2^ and low rhombicity (*R* = 0.07), resulting a significant degree of alignment (1.3 10^−3^). This would subsequently result in ^1^H-^13^C fiRDCs of up to 18 Hz for a completely rigid methyl group.

**Figure 4:**
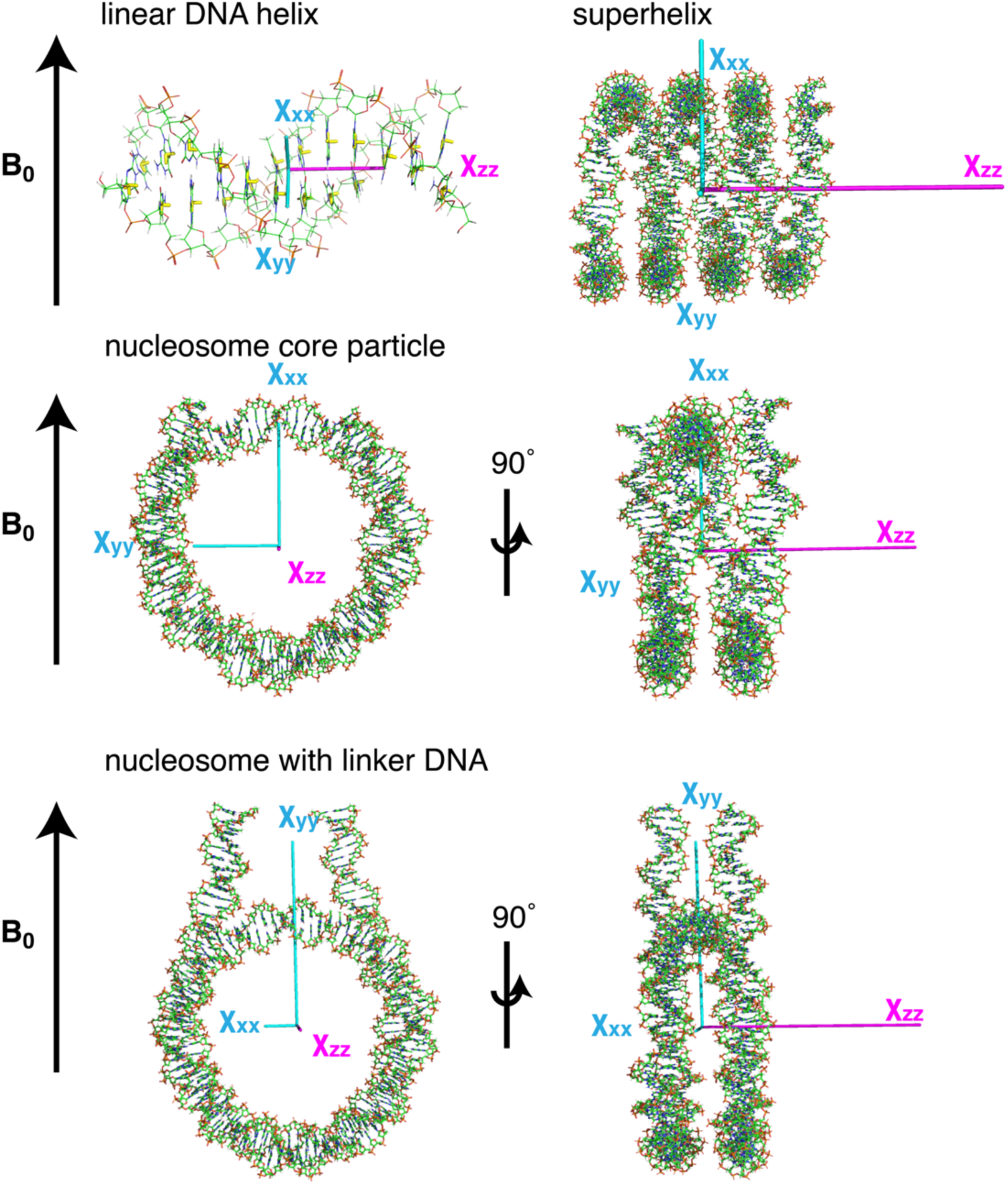
Predicted nucleosome alignment with magnetic field. For linear DNA the largest tensor is predicted along the helix of the DNA. For a superhelix of DNA, the largest tensor aligns along the superhelical axis of the DNA. Both nucleosomes with and without linker DNA have the largest tensor along the superhelical axis, like the superhelix (histone proteins not shown). Linker DNA of 2x 10bp have only a minor influence on the size of the Xzz tensor but have a large influence on the Xxx and Xyy tensor components.

To experimentally determine the extent of magnetic field induced alignment of the nucleosome we recorded IPAP methyl-TROSY experiments^43^ at 600, 900 and 1200 MHz. While at 600 MHz already small deviations from the expected 125 Hz ^1^*J*_CH_ splitting are observed, this becomes particularly clear at 1200 MHz where the splitting ranges between 118.4 and 135.0 Hz (Fig. 5a). This indicates that there is a substantial contribution of fiRDC to the peak splitting, and thus magnetic field induced alignment of the nucleosome. For 33 out of 36 methyl group signals, peak splittings could reliable be measured at all three fields and fitted to extract the ^1^H-^13^C methyl fiRDC at 1200 MHz (Fig. 5b, c). These fiRDCs range from -7.8 to +7.9 Hz. These values are ca. 44% of the predicted maximum fiRDC for the nucleosome core particle. The prediction did however not consider the presence of the additional linker DNA in our system, nor the presence of methyl group dynamics which will downscale the RDC by the order parameter *S*. From the distribution of the fiRDCs, we can estimate that the alignment tensor has high rhombicity (Fig. 5d), which is in sharp contrast with predicted tensor based on the nucleosome core particle structure. This difference in rhombicity is caused by the contribution of the linker DNA and, as we will show below, reflects the conformation of linker DNA.

**Figure 5:**
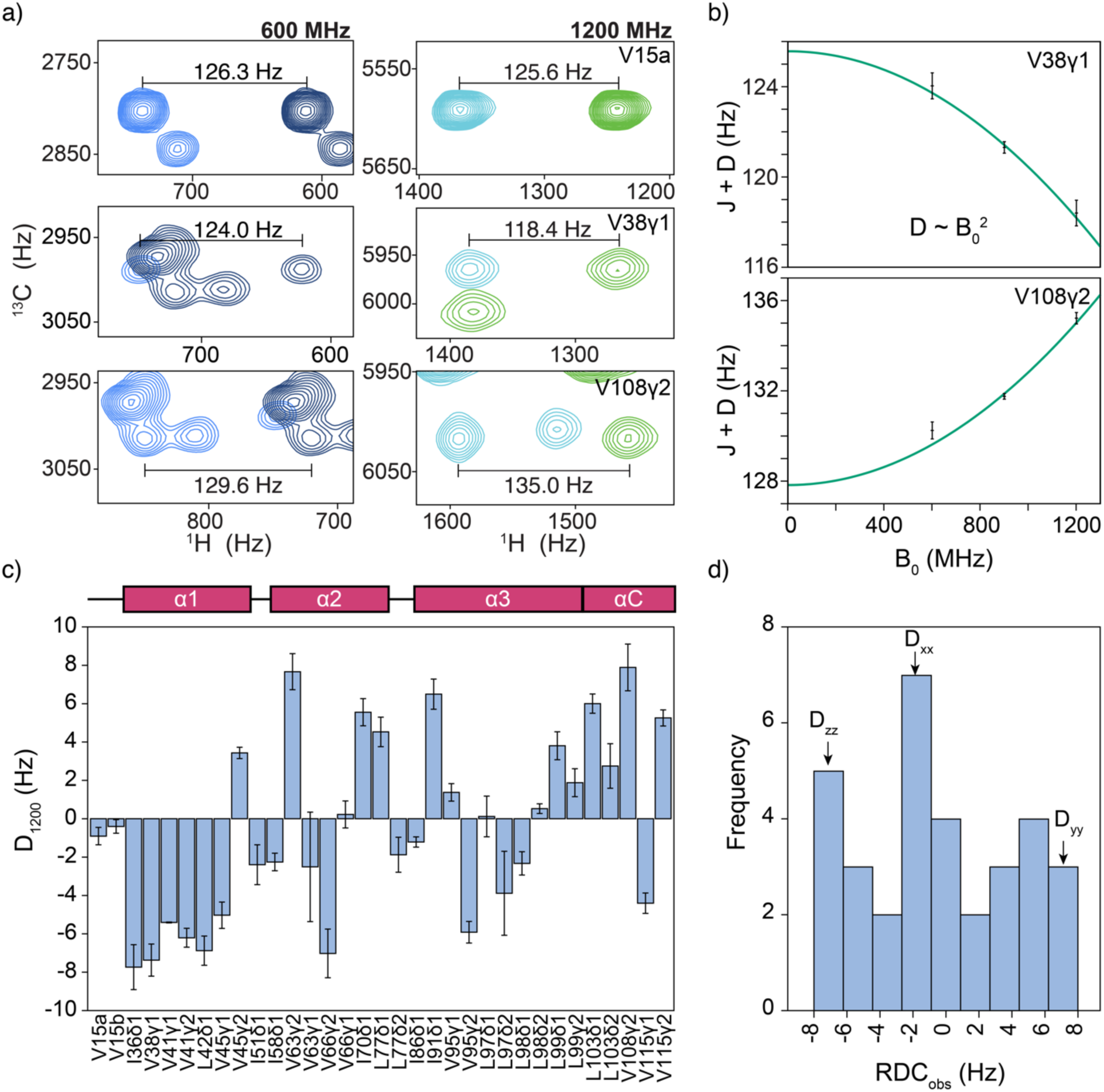
Substantial magnetic field induced alignment of nucleosomes at 28.2 T. a) Zooms of IPAP methyl-TROSY spectra recorded at 600 (14.1 T) and 1200 MHz (28.2 T) showing the ^1^H-^13^C J+D splitting. The spectra with either upfield or downfield doublet component are shown in different colors. Size of the splitting is indicated. b) Experimental field-dependent J+D values and best fits for two selected residues. Full results are in Table S2 and S3. c) Extracted ^1^H-^13^C fiRDC at 1200 MHz for H2B ILV methyl groups in the nucleosome. The secondary structure of H2B is indicated in top of the figure. d) Histogram of the observed fiRDCs at 1200 MHz. The distribution suggests a relatively high rhombicity of the alignment tensor.

### Experimental fiRDCs match very well to nucleosomal H2B structure

We next evaluated the correspondence between the measured fiRDCs and the H2B structure as present in three representative nucleosome structures that have *Xl.* histones. We selected the structure with highest resolution (1KX5) and two high resolution structures that have the non-palindromic 601 DNA sequence that we also use, one determined by crystallography (3LZ0) and one by cryo-electron microscopy (cryo-EM) (7OHC). For each nucleosome structure, the fiRDCs for methyl groups in the histone core with a stereospecific assignment (20 in total) were fitted to the structure of one copy of H2B. The correlation plots of predicted *vs.* observed RDCs show that the best fits are obtained for the structures containing 601 DNA (Fig. 6a). Closer inspection shows that except for three outliers (I58 and L98 methyls), the best correlation is obtained using the 601-nucleosome structure determined by cryo-EM. This suggests that the conformation of I58 and L98 are different in solution from that in the structure, which at least for L98 correlates with its variable conformation in different nucleosome structures (Fig. 3b). Excluding these two residues from the fit resulted in a major improvement in correlation for all structures and overall best result (Q=0.166) for the 1KX5 H2B structure (Fig. 6b).

**Figure 6:**
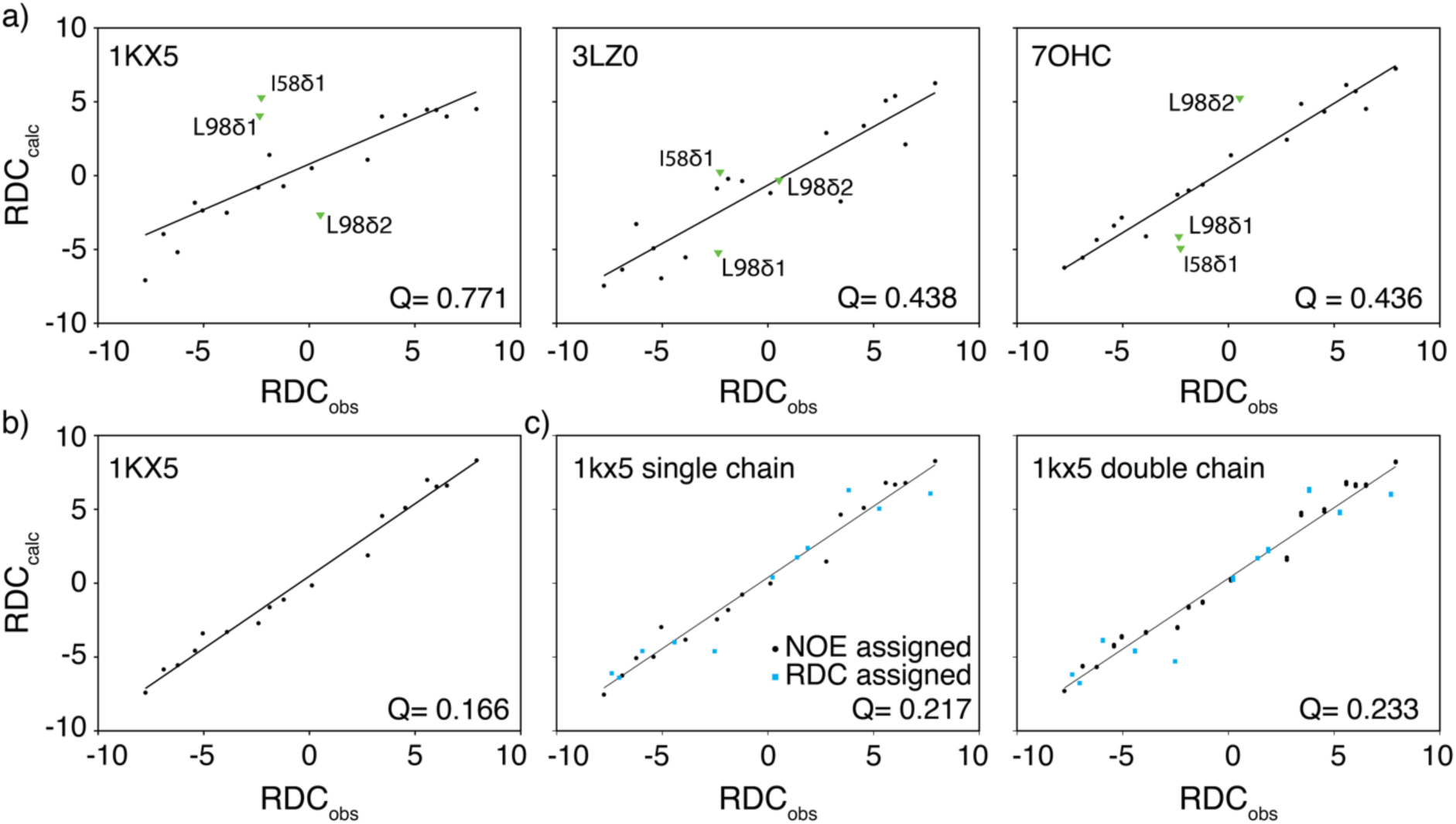
Correlation between experimentally observed and calculated RDCs. RDCs were fit using the indicated structures of H2B in the nucleosome. a) Correlation plots for three structures based on all stereo-specifically assigned residues using NOE data. RDCs for residues I58 and L98 are highlighted. b) Correlation plot excluding I58 and L98 from the fit improves correlation significantly, indicating the solution rotamers for I58 and L98 are different from that in the crystal. c) A correlation plot using both NOE and RDC assigned resonances calculated using a single H2B chain from 1kx5 (left) or both H2B chains (right) show a good correlation (I58 and L98 excluded).

Sidechain rotamer orientation can be determined using residual dipolar couplings^55^. We therefore tested various rotamer conformations of I58 and L98, with all tested rotamers improving the Q-factor compared to 1kx5 rotamer conformations (Table S4 and Fig. S5). For I58 the best fitting rotamer (Table S4 rotamer 1) resulted in a Q-factor (Q=0.165) comparable to the fit excluding I58 and L98. We suggest that I58 adopts this rotamer conformation in solution. For L98 none of tested rotamers led to a satisfactory improvement in Q-factor compared to the fit excluding I58 and L98.

### Determination of stereospecific assignments from fiRDCs

We then reasoned that the predicted RDCs could be used to obtain the stereospecificity of the 6 LV residues (12 methyl groups) for which we could measure a fiRDC but did not have stereospecific assignments. Comparing the experimental fiRDC to the predicted RDC for either the δ1/γ1 or δ2/γ2 methyl group, we could obtain stereospecific assignments for all 6 residues (Table S7). Fitting all 29 fiRDCs from methyl with stereospecific assignment (but excluding I58 and L98) to the 1KX5 H2B structure resulted in an excellent correlation (Q = 0.217) (Fig. 6c). We also tested the feasibility of assigning the stereochemistry if no previous stereochemistry assignments were available as might be the case for other proteins.

Assuming only the stereospecificity of the Ile methyl is known (based on the labeling scheme), 12 out of 14 LV residues could be stereo-specifically assigned after fitting the Ile fiRDCs to the H2B structure (I58 excluded, 5 fiRDCs in total used for the fit). Including chemical shift-based prediction^38^, stereo-specific assignments for 4 Leu could be obtained and in total 12 fiRDCs were used to determine the fit to the H2B structure. Based on the predicted fiRDCs of this fit, again all remaining LV residues could be assigned stereo-specifically, expect for L98 and V41. For L98 neither assignment option gave a good fit, and for V41 both options gave equally good fits. Together, these results show that fiRDCs at ultra-high field are very valuable to aid NOE and chemical shift-based stereo-specific assignment and are potentially also a useful source of information for automated assignment programs.

### Estimation of linker DNA conformation from fiRDCs

Having established that the experimental fiRDCs agree well to the H2B structure, we next wondered whether the fiRDCs could also be used to determine the linker DNA conformation. Linker DNA has been observed with DNA ends bound to the core, detached from the core, or transiently wrap and unwrap from the nucleosome core^56–60^. Additionally, opening of the linkers has been described to occur symmetrically and non-symmetrically^61–63^. The (average) linker DNA conformation in solution can be derived from the fiRDCs, as the extracted alignment tensor can be directly converted into the magnetic susceptibility tensor, which is turn determined mostly by the DNA conformation in the nucleosome. We constructed models of nucleosome with linker DNA in closed or various open states and calculated the Δχ_mol_ from these structures. While the anisotropy of the Δχ_mol_ tensor is not much affected by the conformation of the linker DNA, it is clear from the models that there is strong impact on the rhombicity of the tensor (Fig. 7a and Table S6). The rhombicity is predicted to be highest when the linker DNA extend parallel in the same direction from the nucleosome core particle and decreases upon opening or closing the DNA (Fig. 7a). Thus, comparison of experimental and calculated rhombicity should give a good indication of the average linker DNA conformation. To determine the Δχ_mol_ tensor from the experimental fiRDCs we first extended our analysis from using a single H2B copy to both H2B copies in the nucleosome. Fits to either copy are of similar quality and result in similar orientation of the alignment tensor principal axis frame (PAS) with respect to the molecular frame (Fig. 7b). Indeed, the observation of a single fiRDC for each H2B methyl indicates that the PAS is oriented along the symmetry axis of the nucleosome, as also predicted from the calculated Δχ_mol_ based on the structure (Fig. 7a). Fitting the experimental fiRDCs to both symmetry related H2B copies simultaneously results in a good correlation between observed and predicted RDCs, although Q factor is slightly higher than for the single chain fit (Q= 0.233 vs. 0.217). The alignment tensor from this fit was converted into the magnetic susceptibility tensor to compare against the structure predicted Δχ_mol_ tensors (Fig. 7c). The PAS of the experimental tensor is oriented along the dyad, with the Δχ_zz_ component along the super helical axis and the Δχ_xx_ component perpendicular, in good agreement with orientation of the predicted tensors. The fitted tensor anisotropy is about 60% of the calculated value, reflecting the impact of methyl groups dynamics. While DNA dynamics (transient wrapping and unwrapping of the DNA from the histone core) could also scale the alignment tensor and reduce the fiRDCs, this is less likely to be responsible as the different nucleosome models have all similar anisotropies. Furthermore, FRET measurements have indicated that wrapping/unwrapping occurs at millisecond time scale which is too slow to scale the alignment tensor.

**Figure 7:**
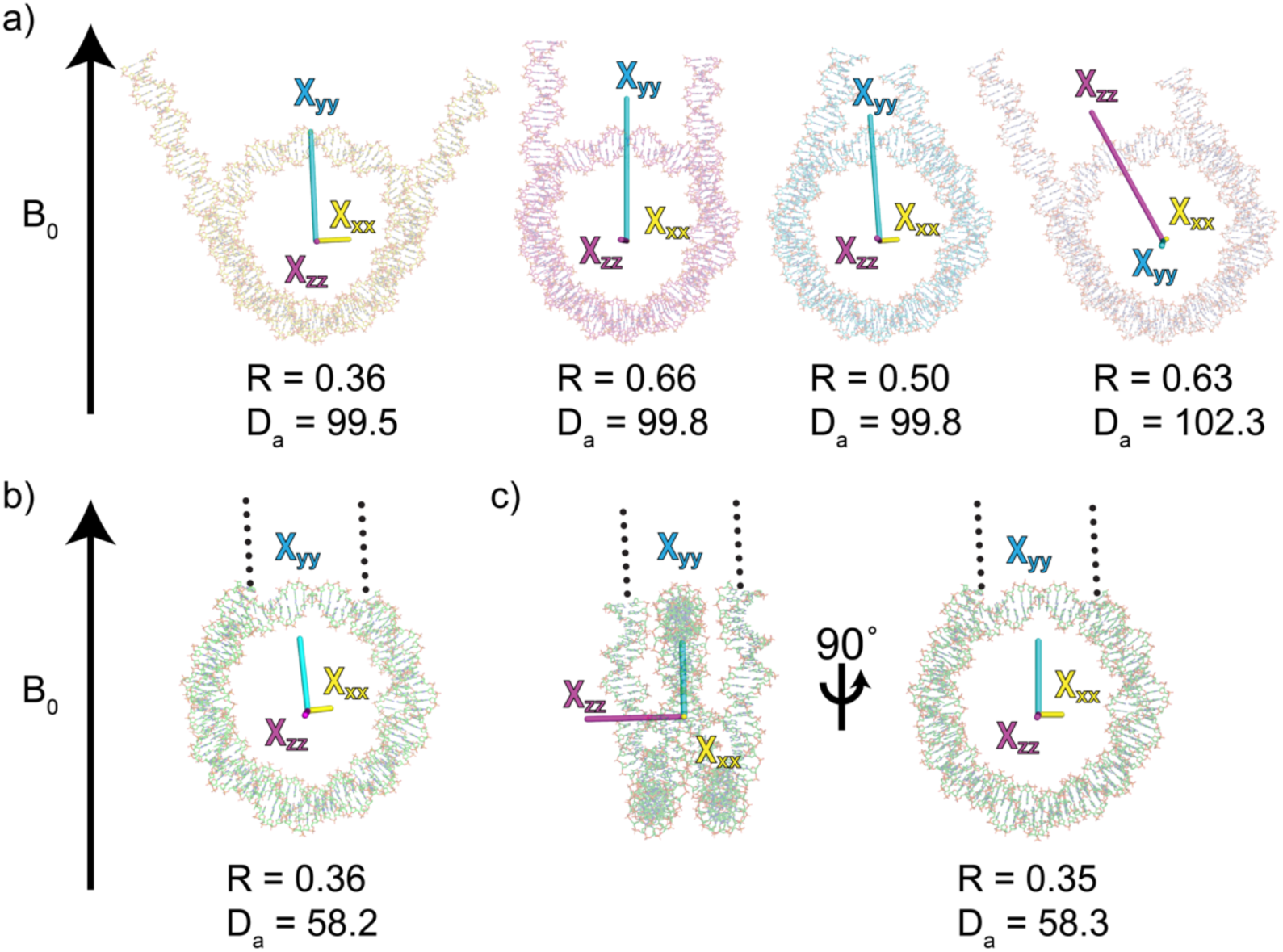
Experimental fiRDCs match an open DNA linker conformation. a) Predicted magnetic susceptibility tensors for 3lz0 with differently modeled linker DNA. From left to right: open, straight, closed, and asymmetric linker DNA. b, c) Observed magnetic susceptibility tensor from the single (b) and double (c) H2B chain fit on 1kx5 nucleosome core particle. Dotted lines represent linker DNA. For each conformation the anisotropy (Da 10^-27^ J T^-2^) is given.

The rhombicity of the Δχ_mol_ tensor could be determined with reasonable precision from the RDC (0.351 ± 0.03) (Fig. 7c). This is substantially lower than the maximal rhombicity for the ‘straight linker’ model and could thus reflect an either more closed or more open structure. As the lowest rhombicity for a compacted linker DNA conformation was predicted to be 0.48, the experimental data suggest that the linker DNA confirmation is rather open. A model with 20 bp unwrapped on both sides from the core particle result in the same rhombicity as experimental (Fig. 7a), indicating that this opened nucleosome represents the time averaged conformation under our conditions.

### Measurable fiRDCs for a small protein-DNA complex

The data on the nucleosome indicate that magnetic field induced alignment at 1200 MHz can make fiRDCs will be an extremely useful tool to study structure and dynamics of large nucleic acid complexes. For protein-only assemblies, or complexes involving shorter nucleic acid fragments, the alignment will likely be significantly smaller. To test the application range of fiRDCs, we finally examined a small protein-DNA complex, formed by the DNA binding domain of pioneer transcription factor Sox2 and a 13 bp double-strand DNA fragment containing its cognate binding site. For a linear B-form DNA of 13 bp, a maximum methyl group RDC of 3.5 Hz is predicted for a bound protein. As Sox2 bends the DNA upon binding, the anisotropies of the nuclesobases will not add up as effectively and a smaller alignment and smaller fiRDCs can be expected. For 8 out 18 LV methyl groups, a fiRDC magnitude larger than 0.2 Hz was obtained, with maximum magnitude of 1.0 Hz, at 1200 MHz (Fig. S6). Overall, the experimental values show reasonable correlation with the structure-based predicted values, albeit that slight differences in DNA and protein conformation give considerable spread in predicted fiRDCs (Fig. S6). Of note, L100 in Sox2 has different side chain conformations in the available structures, resulting in predicted fiRDCs of opposite sign. Experimentally, L100 methyls have close to 0 fiRDCs, suggesting that these different conformations are dynamically averaged in solution (Fig. S6).

## Discussion

We here analyzed the performance of methyl-TROSY NMR at 1.2 GHz with particular attention to use of high resolution and magnetic field induced alignment for the characterization of biomolecular assemblies. We found that for our particular system, the nucleosome, and particular hardware, the methyl-TROSY experiment at 1.2 GHz had on average 20% reduced sensitivity compared to the 900 MHz. It should be noted that the sample had the same concentration at both fields, thus when the same number of spins would be fitted into the active volume of the probe at both fields (3 mm probe and sample at 1200 MHz vs. 5mm probe and sample at 900 MHz), the sensitivity would be increased ∼2.8x at 1200 MHz compared to 900 MHz. The reduced sensitivity is at least in part explained by the increased CSA contribution to the ^13^C transverse relaxation. Our data further indicate that the relative sensitivity of the 1.2 GHz experiment could have been increased somewhat by using slightly longer recycle delays in the fast-pulsing implementation of the methyl-TROSY experiment. Additionally, the use of tailored pulse shapes may increase the relative performance of the 1.2 GHz instrument^66,67,68^.

The increased resolution at 1200 MHz permitted the unambiguous detection of NMR signal duplicities for several histone methyl groups, pointing to a subtle conformational heterogeneity that had been unresolved before. Strikingly, the affected residues correlate with histone side chains that have different conformations between symmetry-related copies in various nucleosome structures. This suggests that the detected conformational heterogeneity may be related to structural differences between symmetry-related histone pairs within the same nucleosome particle.

As a large protein-nucleic acid complex, its DNA component results in a large overall anisotropy in the magnetic susceptibility of the nucleosome, and thus substantial magnetic field induced alignment. Due to the squared dependence on the magnetic field strength, the resulting RDCs are amplified at 1200 MHz. In many cases, such as the Sox2-DNA complex demonstrated here, the field induced RDCs may only become observable at the ultra-high field. For protein-only assemblies, the degree of alignment largely depends on the distribution of aromatic rings in the structure. For symmetric assemblies, the alignment tensors of individual subunits in elongated structures such as the proteasome will add up. For nearly spherical assemblies, such as a viral capsid, the vector-sum will largely cancel out.

The fiRDCs observed for the nucleosome strongly correlated with those predicted for known nucleosome structures and could be used to stereospecifically assign nearly all methyl resonances which were not assignable using conventional NOESY experiments. In cases where a divide-and-conquer approach is not possible or sufficient, assignment of methyl groups in large assemblies is a major challenge, requiring a combination of mutation and NOESY-based strategies. Field-induced alignment is thus a useful addition to obtain stereo-specific assignments and in favorable cases, aid assignment.

Furthermore, the fiRDCs could be used to refine nucleosome structure at both local and global level. At the local level, out of rotamer conformations observed in nucleosome structures, the fiRDCs were used to select the best fitting rotamer for I58. At the global level, we used the fiRDCs to derive that the nucleosome adopts an open conformation of the linker DNA under our conditions. The data indicate that on average not the textbook 1.7, but rather 1.4 superhelical turns of DNA are wrapped around the histone octamer.

Single molecule FRET measurements have shown that presence of linker DNA increases opening of nucleosomes, in particular under low ionic strength conditions such as those used here, suggesting that electrostatic repulsion between the linkers is a dominant factor^64^. Furthermore, the data also indicate that the average linker DNA conformation is symmetric as any asymmetry in linker DNA unwrapping would result in a very different orientation of the PAS (see Fig. 7a). This is in contrast to recent MD study that indicated that linker DNA in 601-nucleosome has intrinsically asymmetric dynamics^63^. It could be that at the longer time scales sampled by the experimental fiRDCs the linkers behave symmetrical, but not at shorter time scale probed by the MD simulations. Alternatively, the asymmetries in DNA dynamics could be too subtle to significantly affect the magnetic susceptibility tensor and thus to be picked up in our experiment.

## Conclusion

Together, our results highlight how ultra-high field methyl-TROSY solution NMR at 1.2 GHz offers greatly improved resolving power and strongly amplified magnetic field induced alignment. Thanks to the ultra-high field data we found evidence for subtle conformational heterogeneities in the nucleosome core and could determine an overall opened structure of the nucleosome in which the DNA is less wrapped compared to crystal or cryo-EM structures. Recently methyltransferases were applied to methylate nucleosomal DNA and used as NMR probes^39^. As nucleosomal DNA is challenging to study by NMR, combination of these type of labeling approaches and fiRDCs can be used to give structural insights in the orientations and dynamics of nucleosomal DNA during critical processes such as nucleosome remodeling and transcription factor binding.

Since field induced alignment is unavoidable as soon as the sample is in the magnet, the combination of ultra-high field and methyl-TROSY solution NMR make field-induced RDCs into a practical tool for the characterization of structural and dynamical features of biomolecular assemblies.

## Acknowledgement

We would like to thank Geeta J. Narlikar (University of California, San Francisco) for providing us with the *Xenopus laevis* nucleosome plasmids. Part of this work was financially supported through iNEXT (PID 5571), provided under the framework of the Horizon 2020 Programme of the European Union through iNEXT (grant 653706) and iNEXT-Discovery (grant 871037).

## Supplemental information

**Figure S1:**
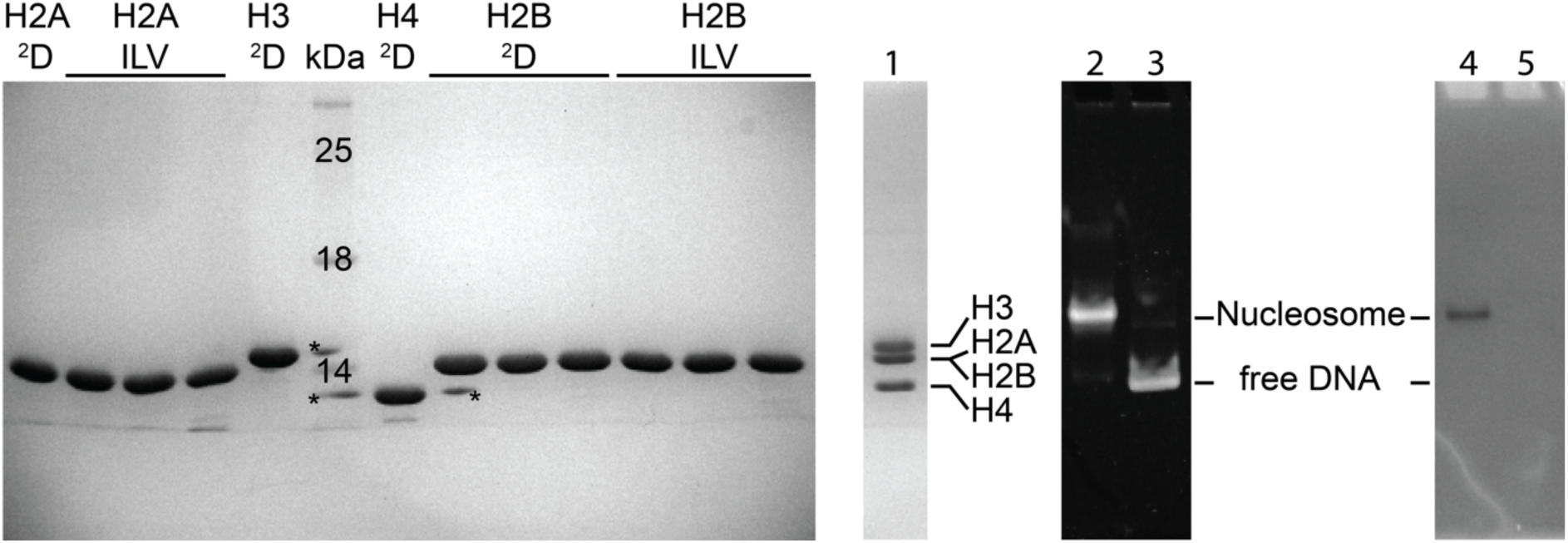
Nucleosome reconstitution. Purified histones (left gel), with stars indicating overflow from neighboring well. 15% SDS page nucleosome (1), 5% native GelRed stain of H2B ILV labeled nucleosome (2) and 167 base pair 601 DNA (3), coomassie stain H2B ILV labeled nucleosome (4) and 167 base pair 601 DNA (5).

**Figure S2:**
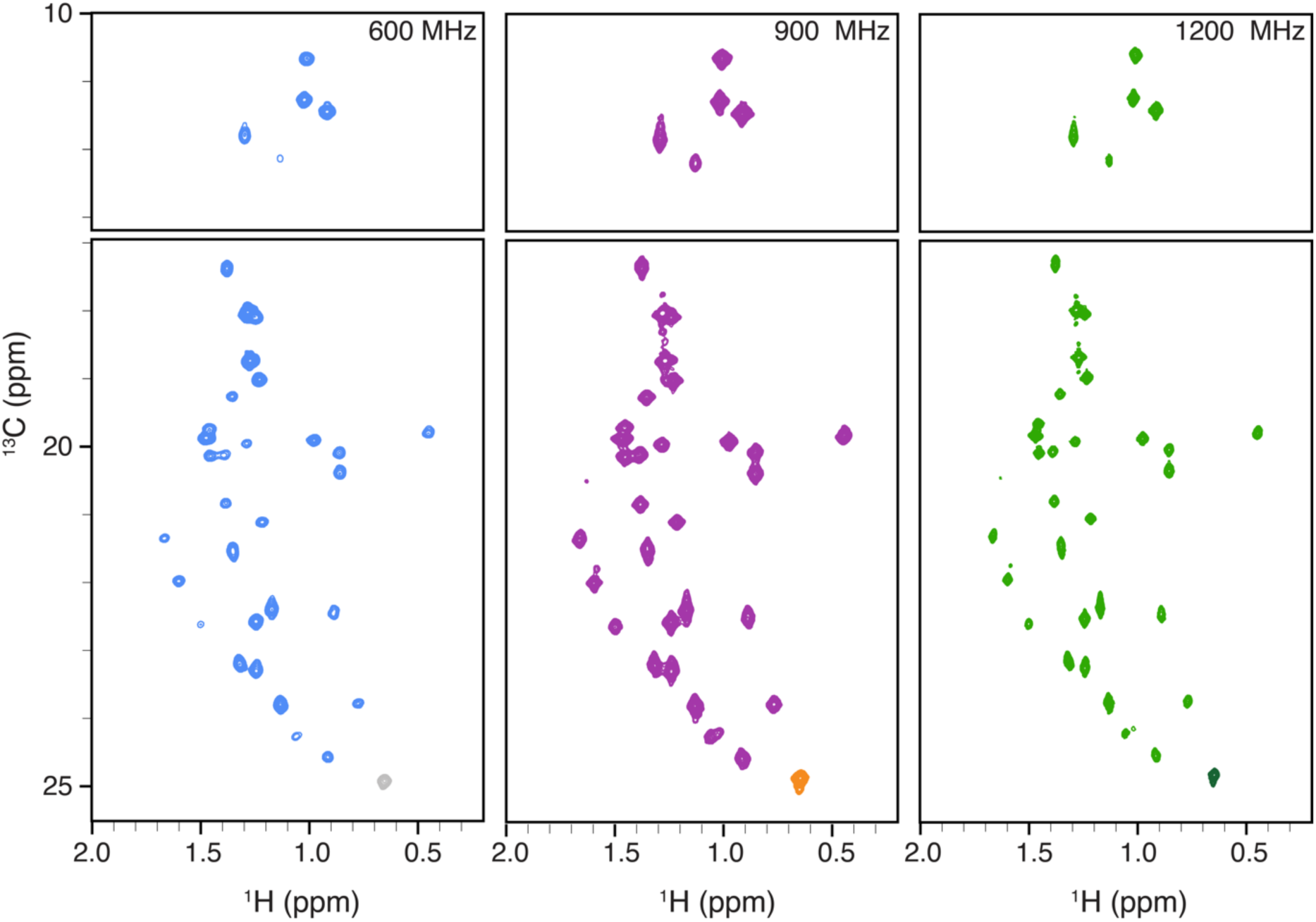
H2B ILV methyl resonances. Nucleosome spectra recorded with SOFAST-methyl TROSY HMQC experiments at 600, 900 and 1200 MHz. Signal threshold is set to compare intensities between different field strengths.

**Figure S3:**
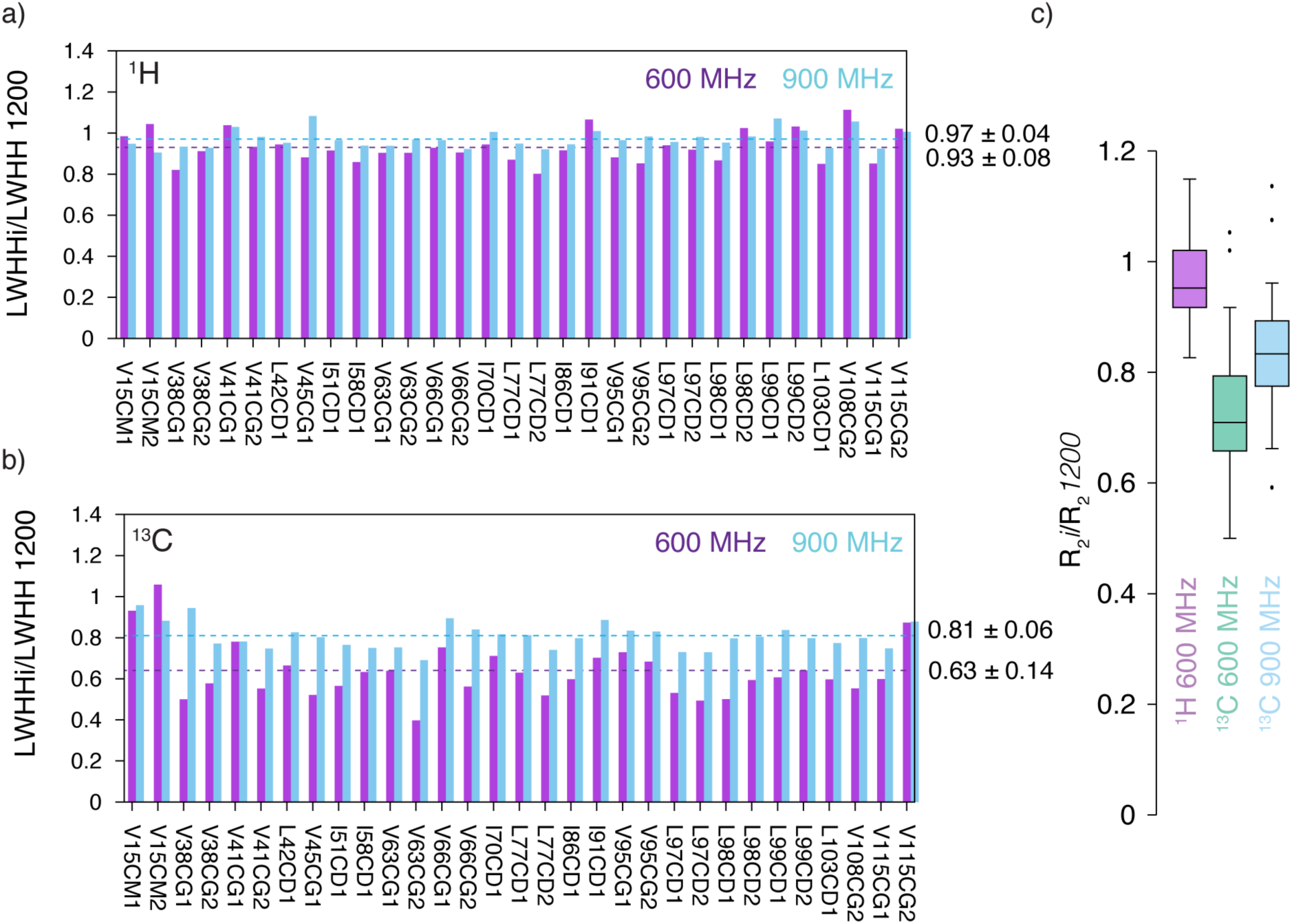
Comparison of ^1^H and ^13^C line widths at 600, 900 and 1200 MHz. Ratio of 1H (a) and 13C (b) line widths at 600 vs. 12000 (purple) and 900 vs. 1200 (blue) MHz, for all non-overlapping methyl signals in the nucleosome. While ^1^H linewidths are very similar at different magnetic field strengths, ^13^C linewidths increase with increasing magnetic field. Line widths measured in Hz using peakfit software on data recorded with 66 (^1^H) and 44 (^13^C) ms resolution, processed without linear prediction. c) Average ratio of R_2_ at given field strength compared to R_2_ at 1200 MHz.

**Figure S4:**
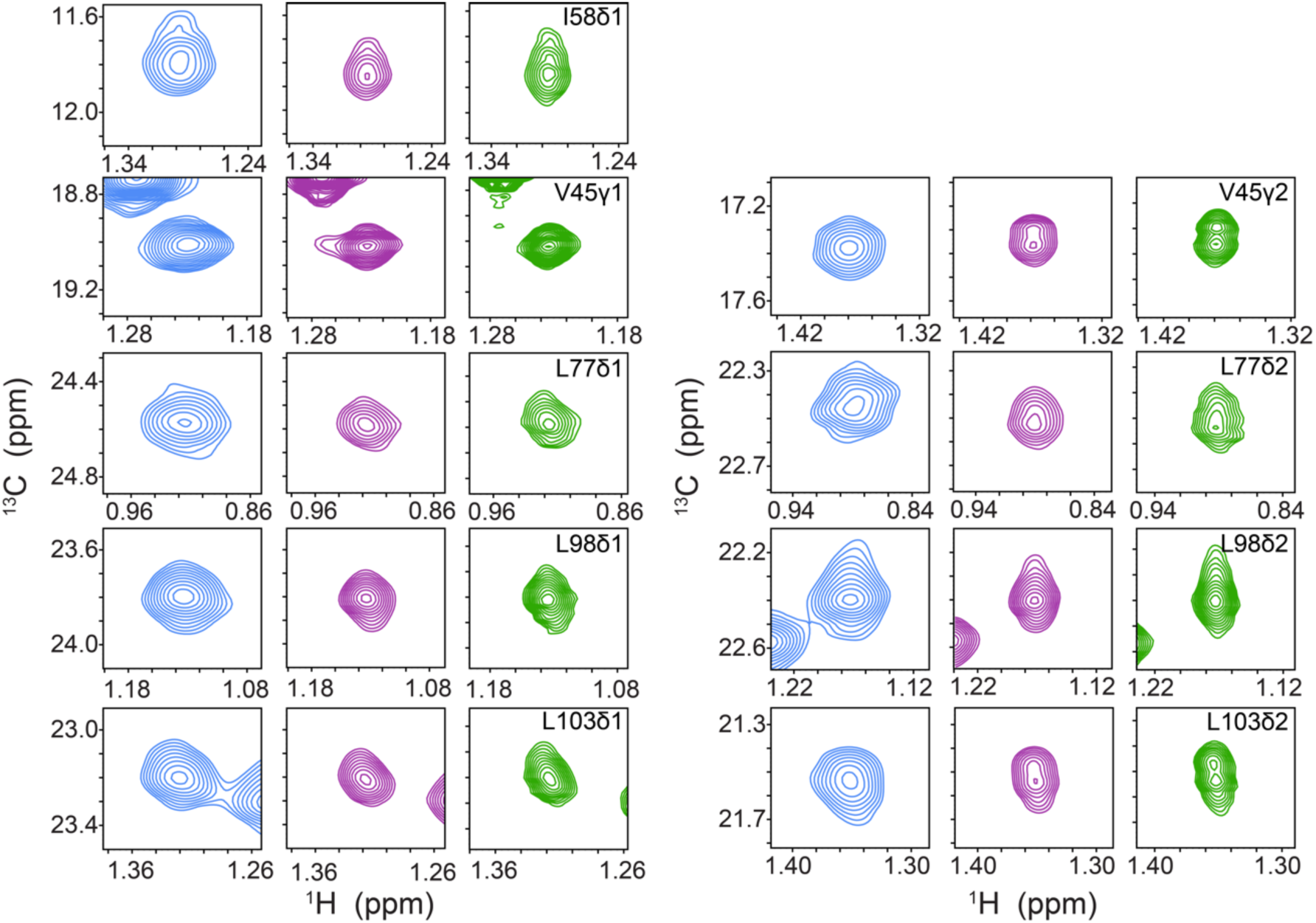
Nucleosome H2B ILV resonance split or broadened peak shapes. Resonances for V45γ2 and L103δ2 show peak doubling. Resonances for I58δ1, L77, L98 and L103δ1 show peak broadening in the ^13^C dimension and L98δ1 and L103δ1 show additional broadening in the ^1^H dimension.

**Figure S5:**
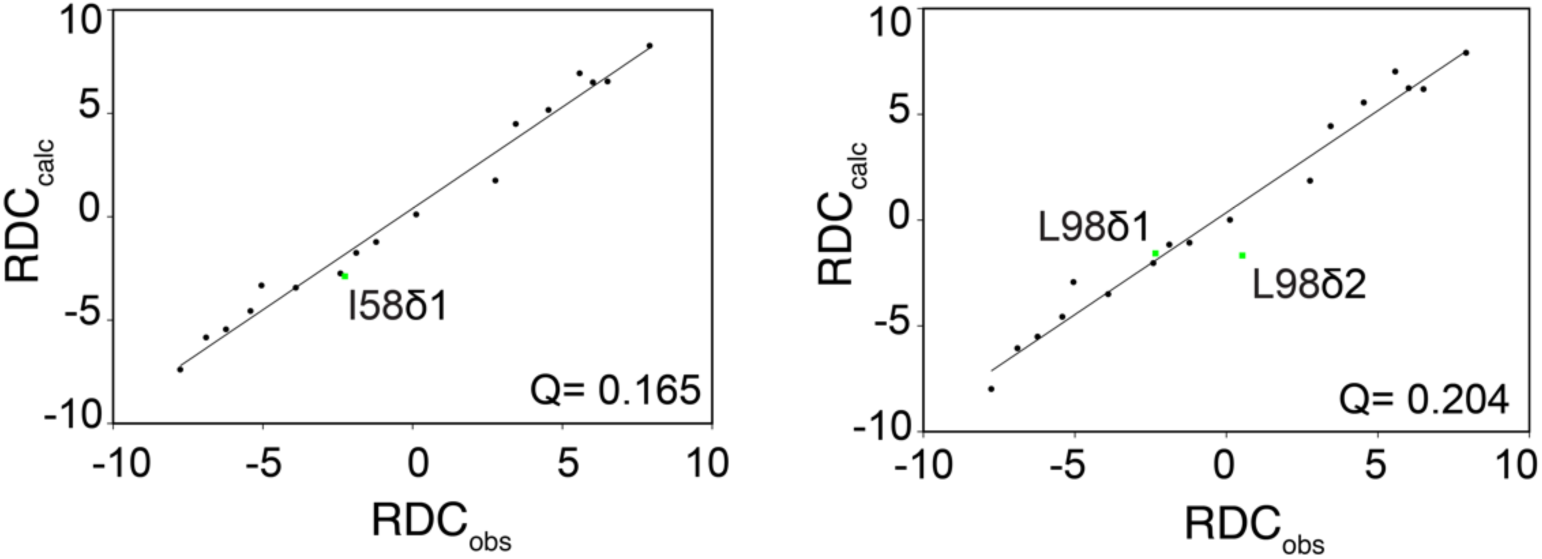
Best fitting alternative rotamer orientations I58 and L98. Residue I58 shows a good correlation using 1kx5 with rotamer 1 for I58 (left). Residue I58 shows an improved correlation using 1kx5 with rotamer 5 for L98 (right).

**Figure S6:**
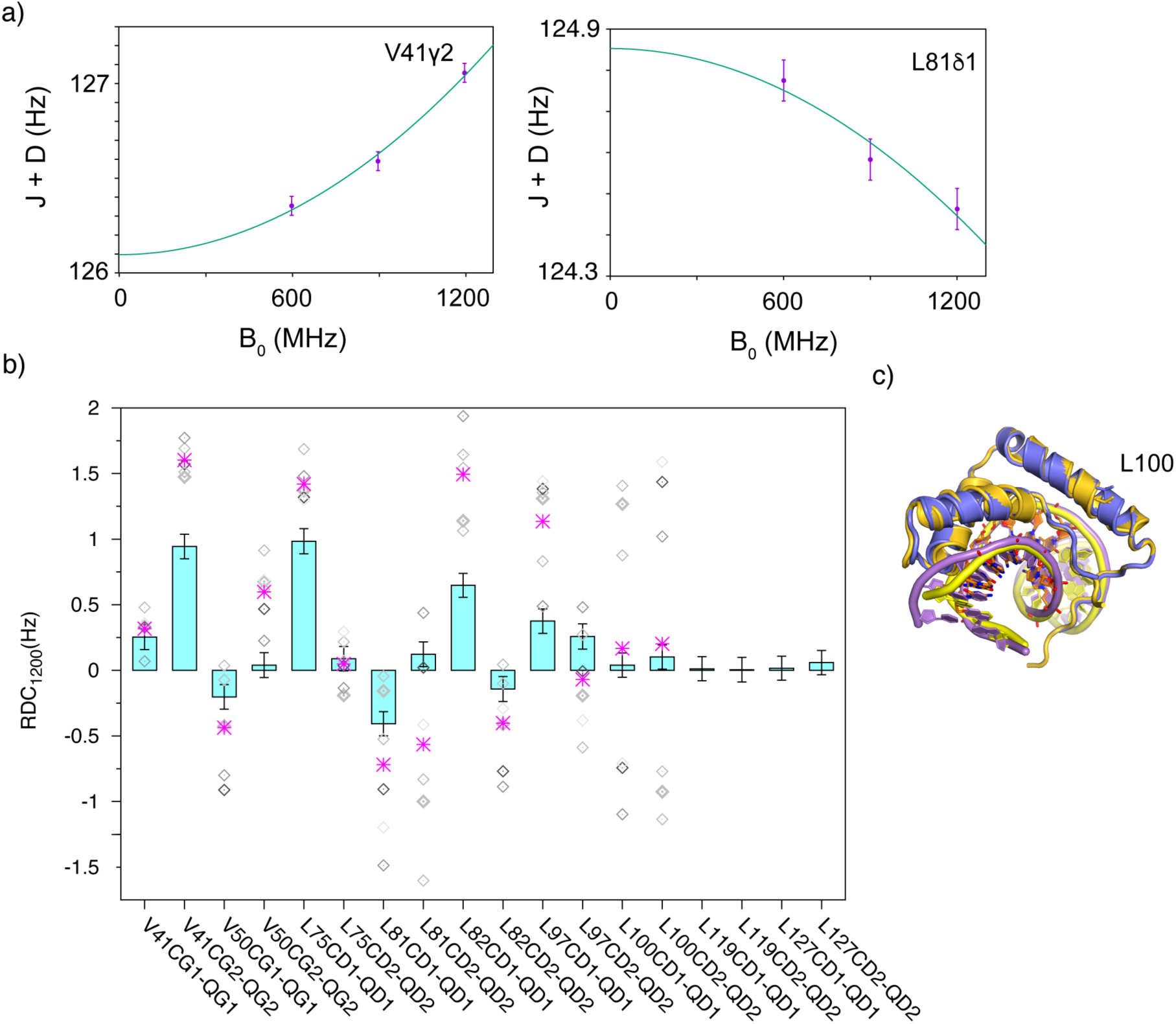
Experimental fiRDCs for Sox2-DBD/DNA complex. (a) Fit of Japp data at 600, 900 and 1200 MHz to extract fiRDCs. (b) Comparison of experimental (blue bars) and structure-predicted (gray symbols) fiRDCs, with average structure-based predicted value in magenta. Structure predicted based on DNA conformation in 1GT0, 6T90 or 6T7B and Sox2 conformation in 1GT0 or 6T90. (c) Overlay of two structures indicating variability in DNA and L100 side chain conformation.

**Table S1:**
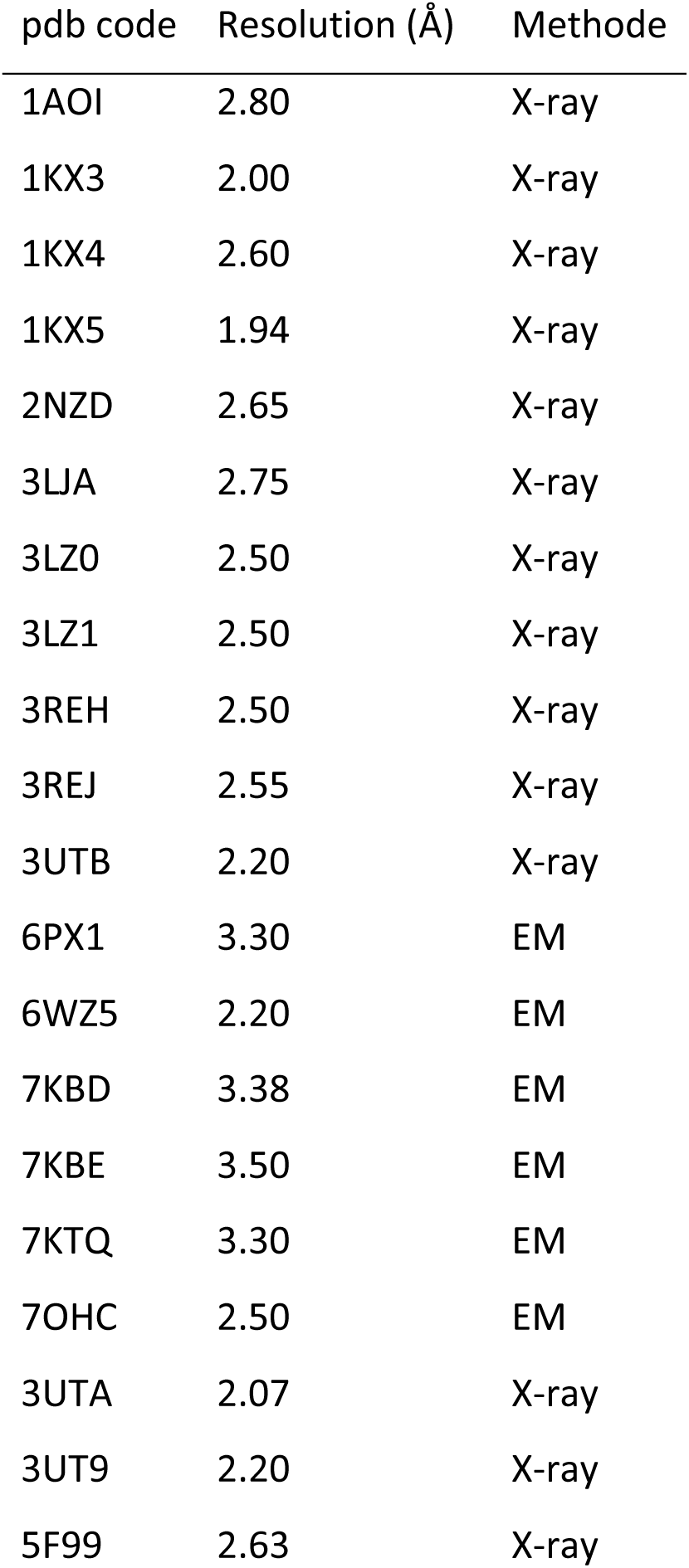
Xenopus laevis Nucleosome pdb structures used for rotamer analysis.

| pdb code | Resolution (Å) | Methode |
| --- | --- | --- |
| 1AOI | 2.80 | X-ray |
| 1KX3 | 2.00 | X-ray |
| 1KX4 | 2.60 | X-ray |
| 1KX5 | 1.94 | X-ray |
| 2NZD | 2.65 | X-ray |
| 3LJA | 2.75 | X-ray |
| 3LZ0 | 2.50 | X-ray |
| 3LZ1 | 2.50 | X-ray |
| 3REH | 2.50 | X-ray |
| 3REJ | 2.55 | X-ray |
| 3UTB | 2.20 | X-ray |
| 6PX1 | 3.30 | EM |
| 6WZ5 | 2.20 | EM |
| 7KBD | 3.38 | EM |
| 7KBE | 3.50 | EM |
| 7KTQ | 3.30 | EM |
| 7OHC | 2.50 | EM |
| 3UTA | 2.07 | X-ray |
| 3UT9 | 2.20 | X-ray |
| 5F99 | 2.63 | X-ray |

**Table S2:** Carbon coupled peak splitting. Peak splitting (Δ) at 600, 900 and 1200 MHz due to carbon coupling are reported in Hz for nucleosome H2B ILV methyl groups.

| residue | methyl | $\Delta_{600}$ | $\sigma_{S,600}$ | $\Delta_{900}$ | $\sigma_{S,900}$ | $\Delta_{1200}$ | $\sigma_{S,1200}$ |
| --- | --- | --- | --- | --- | --- | --- | --- |
| V15 | a | 126.55 | 0.05 | 125.63 | 0.01 | 125.34 | 0.02 |
| V15 | b | 126.26 | 0.04 | 125.84 | 0.01 | 125.74 | 0.02 |
| I36 | $\delta 1$ | 125.45 | 0.79 | 122.25 | 0.24 | 119.19 | 0.56 |
| V38 | $\gamma 1$ | 124.28 | 0.58 | 121.31 | 0.25 | 118.40 | 0.57 |
| V41 | $\gamma 1$ | 125.70 | 0.39 | 123.87 | 0.15 | 121.49 | 0.29 |
| V41 | $\gamma 2$ | 124.94 | 0.43 | 122.84 | 0.16 | 120.28 | 0.35 |
| L42 | $\delta 1$ | 123.49 | 0.39 | 120.86 | 0.14 | 118.07 | 0.27 |
| V45 | $\gamma 1$ | 125.91 | 0.14 | 123.56 | 0.05 | 121.55 | 0.09 |
| V45 | $\gamma 2$ | 126.50 | 0.21 | 127.62 | 0.08 | 129.23 | 0.19 |
| I51 | $\delta 1$ | 125.24 | 0.27 | 123.75 | 0.10 | 123.03 | 0.21 |
| I58 | $\delta 1$ | 125.70 | 0.27 | 124.89 | 0.12 | 124.08 | 0.27 |
| V63 | $\gamma 2$ | 127.19 | 0.55 | 130.28 | 0.23 | 133.29 | 0.53 |
| V63 | $\gamma 1$ | 125.89 | 0.43 | 122.80 | 0.18 | 120.62 | 0.42 |
| V66 | $\gamma 2$ | 125.70 | 0.30 | 122.77 | 0.12 | 120.17 | 0.30 |
| V66 | $\gamma 1$ | 126.43 | 0.27 | 125.93 | 0.09 | 126.23 | 0.17 |
| I70 | $\delta 1$ | 127.02 | 0.22 | 128.28 | 0.08 | 130.93 | 0.15 |
| L77 | $\delta 1$ | 125.77 | 0.41 | 126.71 | 0.18 | 129.00 | 0.43 |
| L77 | $\delta 2$ | 124.53 | 0.39 | 123.70 | 0.17 | 123.25 | 0.43 |
| I86 | $\delta 1$ | 125.35 | 0.17 | 124.72 | 0.06 | 124.27 | 0.12 |
| I91 | $\delta 1$ | 127.51 | 0.29 | 128.75 | 0.10 | 131.83 | 0.19 |
| V95 | $\gamma 1$ | 126.75 | 0.45 | 126.99 | 0.15 | 127.72 | 0.28 |
| V95 | $\gamma 2$ | 125.27 | 0.69 | 123.75 | 0.24 | 120.98 | 0.52 |
| L97 | $\delta 1$ | 125.17 | 0.46 | 124.80 | 0.20 | 125.26 | 0.46 |
| L97 | $\delta 2$ | 124.57 | 0.38 | 122.24 | 0.15 | 121.26 | 0.30 |
| L98 | $\delta 1$ | 124.22 | 0.15 | 123.37 | 0.06 | 122.55 | 0.12 |
| L98 | $\delta 2$ | 124.82 | 0.16 | 124.87 | 0.06 | 125.18 | 0.12 |
| L99 | $\delta 1$ | 125.64 | 0.24 | 126.53 | 0.08 | 128.41 | 0.16 |
| L99 | $\delta 2$ | 124.87 | 0.17 | 125.23 | 0.06 | 126.27 | 0.12 |
| L103 | $\delta 1$ | 126.21 | 0.22 | 127.86 | 0.08 | 130.66 | 0.17 |
| L103 | $\delta 2$ | 126.26 | 0.22 | 126.63 | 0.08 | 128.21 | 0.18 |
| V108 | $\gamma 2$ | 129.45 | 0.41 | 131.23 | 0.14 | 135.04 | 0.28 |
| V115 | $\gamma 1$ | 125.81 | 0.16 | 124.18 | 0.05 | 122.39 | 0.10 |
| V115 | $\gamma 2$ | 127.39 | 0.27 | 128.90 | 0.08 | 131.31 | 0.14 |

**Table S3:** Fitting of nucleosome RDC at 1200 MHz. J_CH_ and RDC of H2B ILV labeled nucleosomes are fit with fit parameters and errors reported in Hz.

| residue | methyl | $J_{CH}$ | $\sigma_{JCH}$ | $c_{rdc}(10^{-6})$ | $\sigma_{crdc}(10^{-6})$ | fiRDC | $\sigma_{fiRDC}$ |
| --- | --- | --- | --- | --- | --- | --- | --- |
| V15 | a | 126.20 | 0.32 | -0.63 | 0.32 | -0.91 | 0.46 |
| V15 | b | 126.10 | 0.25 | -0.28 | 0.25 | -0.40 | 0.35 |
| I36 | $\delta 1$ | 126.72 | 0.74 | -5.39 | 0.82 | -7.76 | 1.18 |
| V38 | $\gamma 1$ | 125.57 | 0.52 | -5.13 | 0.58 | -7.39 | 0.84 |
| V41 | $\gamma 1$ | 126.92 | 0.02 | -3.77 | 0.02 | -5.42 | 0.02 |
| V41 | $\gamma 2$ | 126.40 | 0.31 | -4.32 | 0.34 | -6.23 | 0.49 |
| L42 | $\delta 1$ | 124.83 | 0.50 | -4.79 | 0.53 | -6.90 | 0.76 |
| V45 | $\gamma 1$ | 126.48 | 0.46 | -3.50 | 0.48 | -5.05 | 0.69 |
| V45 | $\gamma 2$ | 125.71 | 0.18 | 2.39 | 0.21 | 3.44 | 0.30 |
| I51 | $\delta 1$ | 125.22 | 0.67 | -1.67 | 0.72 | -2.41 | 1.04 |
| I58 | $\delta 1$ | 126.22 | 0.28 | -1.57 | 0.32 | -2.26 | 0.46 |
| V63 | $\gamma 2$ | 125.84 | 0.58 | 5.34 | 0.65 | 7.69 | 0.94 |
| V63 | $\gamma 1$ | 123.89 | 1.75 | -1.75 | 1.99 | -2.52 | 2.86 |
| V66 | $\gamma 2$ | 126.87 | 0.77 | -4.89 | 0.88 | -7.04 | 1.27 |
| V66 | $\gamma 1$ | 125.89 | 0.47 | 0.15 | 0.49 | 0.22 | 0.71 |
| I70 | $\delta 1$ | 125.23 | 0.47 | 3.87 | 0.49 | 5.57 | 0.71 |
| L77 | $\delta 1$ | 124.24 | 0.47 | 3.15 | 0.54 | 4.54 | 0.77 |
| L77 | $\delta 2$ | 124.87 | 0.55 | -1.31 | 0.64 | -1.88 | 0.92 |
| I86 | $\delta 1$ | 125.44 | 0.17 | -0.84 | 0.19 | -1.22 | 0.27 |
| I91 | $\delta 1$ | 125.18 | 0.52 | 4.52 | 0.55 | 6.51 | 0.79 |
| V95 | $\gamma 1$ | 126.27 | 0.30 | 0.96 | 0.32 | 1.38 | 0.46 |
| V95 | $\gamma 2$ | 127.03 | 0.36 | -4.12 | 0.39 | -5.94 | 0.56 |
| L97 | $\delta 1$ | 124.86 | 0.65 | 0.08 | 0.74 | 0.12 | 1.06 |
| L97 | $\delta 2$ | 124.70 | 1.40 | -2.71 | 1.52 | -3.90 | 2.19 |
| L98 | $\delta 1$ | 124.75 | 0.38 | -1.62 | 0.42 | -2.34 | 0.61 |
| L98 | $\delta 2$ | 124.60 | 0.16 | 0.37 | 0.18 | 0.53 | 0.25 |
| L99 | $\delta 1$ | 124.47 | 0.48 | 2.65 | 0.51 | 3.82 | 0.73 |
| L99 | $\delta 2$ | 124.26 | 0.48 | 1.31 | 0.51 | 1.88 | 0.73 |
| L103 | $\delta 1$ | 124.53 | 0.32 | 4.18 | 0.35 | 6.02 | 0.51 |
| L103 | $\delta 2$ | 125.20 | 0.74 | 1.91 | 0.81 | 2.76 | 1.17 |
| V108 | $\gamma 2$ | 126.91 | 0.79 | 5.49 | 0.85 | 7.91 | 1.22 |
| V115 | $\gamma 1$ | 126.72 | 0.35 | -3.07 | 0.37 | -4.42 | 0.53 |
| V115 | $\gamma 2$ | 125.98 | 0.28 | 3.66 | 0.29 | 5.27 | 0.42 |

**Table S4:**
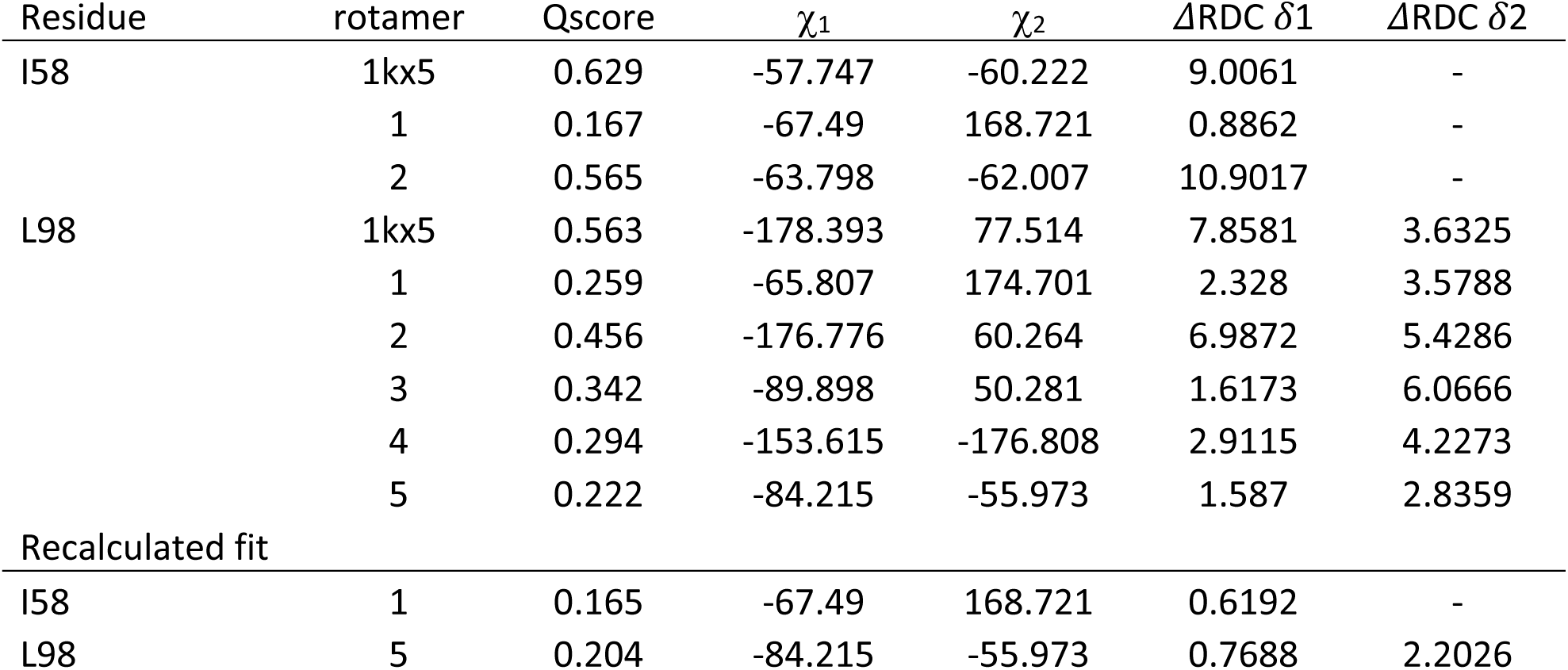
Alternative rotamer scores I58 and L98. Q-scores calculated for 1kx5 and alternative rotamer conformations for residue I58 and L98. Top part contains RDCs for I58 and L98 where I58 and L98 are not used to determine the fit. Bottom part contains RDCs where I58 or L98 are used as input for the fitting. Lower Q-scores indicate a better fit for rotamers with rotation angles chi1 and chi2. Difference between predicted and observed RDCs is given by *Δ*RDC *δ*1 and *Δ*RDC *δ*2 for the *δ*1 and *δ*2 methyl groups. Residue I58 has a low *Δ*RDC for rotamer 1 indicating a good rotamer orientation. Residue L98 has a low *Δ*RDC *δ*1 but high *Δ*RDC *δ*2 for rotamer 5 indicating that this rotamer is not yet optimal.

**Table S5:** Stereochemistry assignments based on RDC, Sider and transferred peaks. Final stereochemistry is given together with the observed RDC. Predicted RDC based on a subset of only isoleucine or isoleucine and sider predicted leucine or all known stereochemistry. Input for predictions is indicated in gray. Confidence of stereochemistry assignment is given for less than 2 σ (dark green) and less than 4 σ (light green). When no assignment was possible values are indicated in red for more than 4 σ, or yellow if no assignment because two RDC’s are too close to each other (less than one σ difference).

| residue | methyl | RDC<sub>observed</sub> | $\sigma$ (hz) | RDC<sub>Pred</sub> ile | RDC<sub>Pred</sub> ile+leu | RDC<sub>Pred</sub> transferred |
| --- | --- | --- | --- | --- | --- | --- |
| I36 | $\delta 1$ | -7.76 | 1.18 | -7.76 | -7.37 | -7.42 |
| I51 | $\delta 1$ | -2.41 | 1.04 | -2.42 | -2.22 | -2.72 |
| I58 | $\delta 1$ | -2.26 | 0.46 | 8.35 | 8.88 | 9.26 |
| I70 | $\delta 1$ | 5.57 | 0.71 | 5.57 | 6.65 | 6.99 |
| I86 | $\delta 1$ | -1.22 | 0.27 | -1.21 | -0.76 | -1.12 |
| I91 | $\delta 1$ | 6.51 | 0.79 | 6.50 | 6.44 | 6.61 |
| L42 | $\delta 1$ | -6.90 | 0.76 | -6.71 | -6.09 | -5.83 |
| L42 | $\delta 2$ | - | | 1.82 | 1.76 | 1.78 |
| L77 | $\delta 1$ | 4.54 | 0.77 | 5.26 | 5.07 | 5.11 |
| L77 | $\delta 2$ | -1.88 | 0.92 | -2.33 | -1.72 | -1.63 |
| L97 | $\delta 1$ | 0.12 | 1.06 | 2.82 | -0.05 | -0.14 |
| L97 | $\delta 2$ | -3.90 | 2.19 | -5.83 | -3.69 | -3.30 |
| L98 | $\delta 1$ | -2.34 | 0.61 | 6.84 | 7.52 | 7.84 |
| L98 | $\delta 2$ | 0.53 | 0.25 | -6.44 | -4.97 | -4.58 |
| L99 | $\delta 1$ | 3.82 | 0.73 | 4.76 | 6.12 | 6.51 |
| L99 | $\delta 2$ | 1.88 | 0.73 | 2.20 | 2.24 | 2.22 |
| L103 | $\delta 1$ | 6.02 | 0.51 | 6.70 | 6.35 | 6.54 |
| L103 | $\delta 2$ | 2.76 | 1.17 | -0.77 | 1.50 | 1.89 |
| V38 | $\gamma 1$ | -7.39 | 0.84 | -4.42 | -5.93 | -6.33 |
| V38 | $\gamma 2$ | - | - | 4.40 | 4.61 | 4.68 |
| V41 | $\gamma 1$ | -5.42 | 0.02 | -4.70 | -4.88 | -4.57 |
| V41 | $\gamma 2$ | -6.23 | 0.49 | -3.89 | -4.86 | -5.56 |
| V45 | $\gamma 1$ | -5.05 | 0.69 | -2.08 | -2.76 | -3.40 |
| V45 | $\gamma 2$ | 3.44 | 0.30 | 4.91 | 4.40 | 4.54 |
| V63 | $\gamma 1$ | 7.69 | 0.94 | 7.98 | 5.91 | 5.92 |
| V63 | $\gamma 2$ | -2.52 | 2.86 | -4.38 | -4.45 | -5.10 |
| V66 | $\gamma 1$ | -7.04 | 1.27 | -5.31 | -6.21 | -6.76 |
| V66 | $\gamma 2$ | 0.22 | 0.71 | 0.17 | 0.36 | 0.30 |
| V95 | $\gamma 1$ | 1.38 | 0.46 | 4.41 | 1.72 | 1.51 |
| V95 | $\gamma 2$ | -5.94 | 0.56 | -5.15 | -4.45 | -4.08 |
| V108 | $\gamma 1$ | - | - | -2.90 | -2.63 | -3.18 |
| V108 | $\gamma 2$ | 7.91 | 1.22 | 7.32 | 7.96 | 8.31 |
| V115 | $\gamma 1$ | -4.42 | 0.53 | -2.51 | -3.80 | -4.48 |
| V115 | $\gamma_2$ | 5.27 | 0.42 | 4.45 | 4.77 | 4.83 |

**Table S6:** RDC fit and model scores.

| **Measured** | Q-score | Da ($10^{-4}$) | Dr ($10^{-4}$) | R |
| --- | --- | --- | --- | --- |
| 1xk5 chain D | 0.771 | 0.92 | 0.46 | 0.504 |
| 3lz0 chain H | 0.438 | -1.12 | -0.61 | 0.547 |
| 7ohc chain D | 0.436 | -0.95 | -0.46 | 0.486 |
| 1xk5 -I58,L98 | 0.166 | -1.17 | -0.43 | 0.364 |
| 3lz0 -I58,L98 | 0.414 | -1.15 | -0.67 | 0.579 |
| 7ohc -I58,L98 | 0.203 | -1.09 | -0.42 | 0.386 |
| **assigned** |  |  |  |  |
| 1kx5 single chain | 0.217 | -1.17 | -0.42 | 0.355 |
| 1kx5 both chains | 0.233 | -1.17 | -0.40 | 0.345 |
| **Predicted** |  |  |  |  |
| closed | - | - | - | 0.501 |
| straight | - | - | - | 0.662 |
| open | - | - | - | 0.357 |
| assymetric | - | - | - | 0.630 |

**Table S7:**
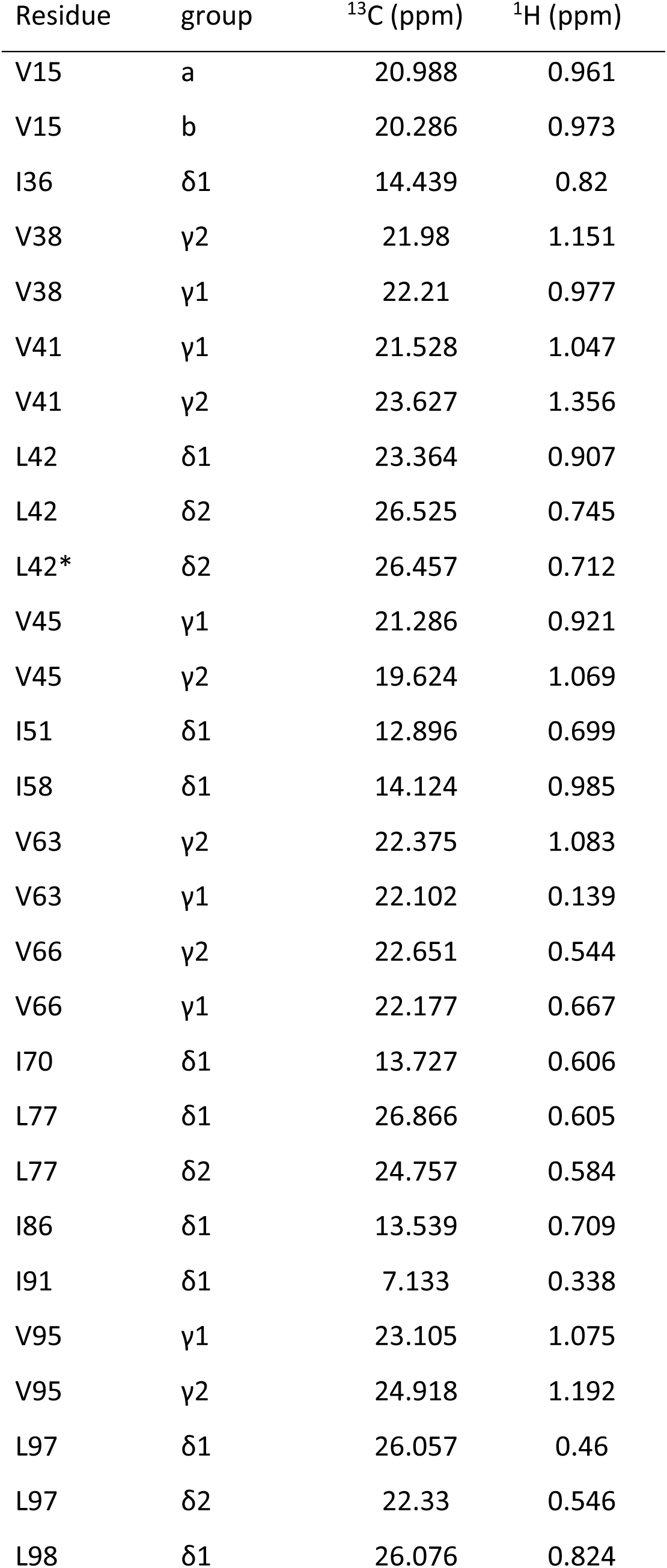
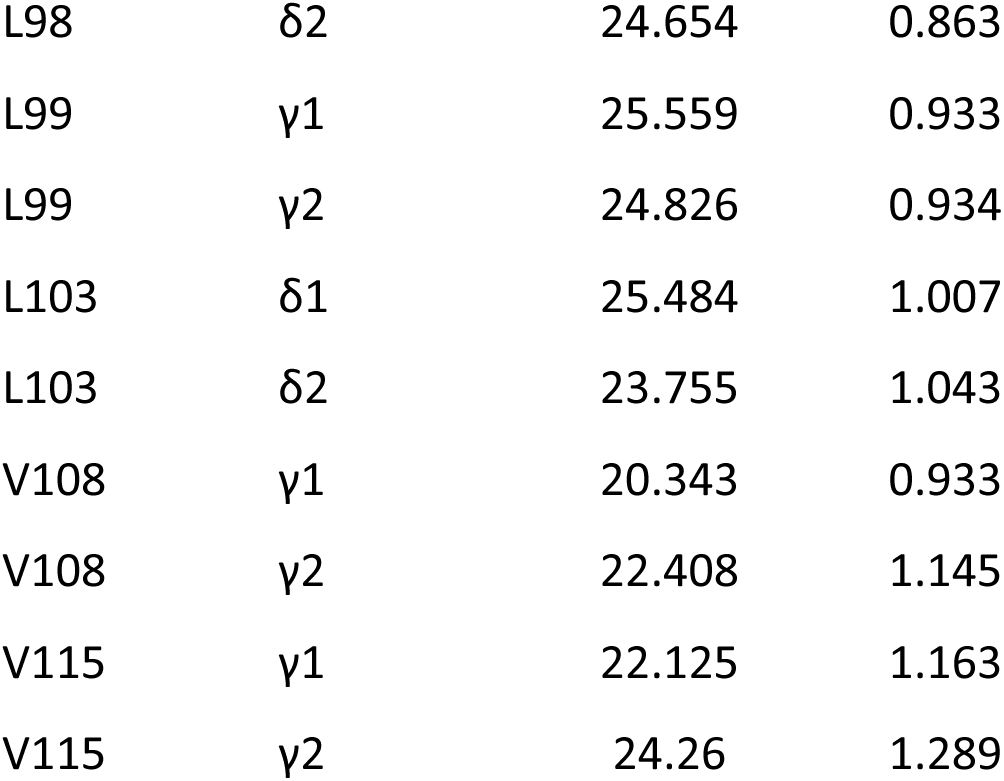
*Xenopus laevis* H2B ILV nucleosome assignment. Nucleosome residue and stereospecific assignment for Xl. ILV labeled nucleosomal H2B. Notations a and b indicate an unknown stereospecific assignment.

